# Fibroblast activation protein regulates natural killer cell migration, extravasation and tumor infiltration

**DOI:** 10.1101/2021.02.03.429622

**Authors:** Allison A. Fitzgerald, Rachael E. Maynard, Emily F. Marcisak, Apsra Nasir, Eric Glasgow, Sandra A. Jablonski, Pieter Van Der Veken, Gray Pearson, Shira E. Eisman, Emily M. Mace, Elana J. Fertig, Louis M. Weiner

**Affiliations:** Department of Oncology, Georgetown Lombardi Comprehensive Cancer Center, Georgetown University Medical Center, Washington, DC, USA; McKusick-Nathans Institute of the Department of Genetic Medicine, Johns Hopkins School of Medicine, Baltimore, MD, USA; Department of Oncology, Sidney Kimmel Comprehensive Cancer Center, Johns Hopkins School of Medicine, Baltimore, MD, USA; Department of Medicinal Chemistry/UAMC, University of Antwerp, Universiteitsplein 1, B2610 Wilrijk, Belgium; Department of Applied Mathematics and Statistics, Johns Hopkins University Whiting School of Engineering, Baltimore, MD, USA; Department of Biomedical Engineering, Johns Hopkins University School of Medicine, Baltimore, MD, USA; Department of Pediatrics, Vagelos College of Physicians and Surgeons, Columbia University Irving Medical Center, New York, NY, USA

## Abstract

Natural killer (NK) cells play a critical role in physiologic and pathologic conditions such as pregnancy, infection, autoimmune disease and cancer. In cancer, numerous strategies have been designed to exploit the cytolytic properties of NK cells, with variable success. A major hurdle to NK-cell focused therapies is NK cell recruitment and infiltration into tumors. While the chemotaxis pathways regulating NK recruitment to different tissues are well delineated, the mechanisms human NK cells employ to physically migrate are ill-defined. We show for the first time that human NK cells express fibroblast activation protein (FAP), a cell surface protease previously thought to be primarily expressed by activated fibroblasts. FAP degrades the extracellular matrix to facilitate cell migration and tissue remodeling. We used novel *in vivo* zebrafish and *in vitro* 3D culture models to demonstrate that FAP knock out and pharmacologic inhibition restrict NK cell migration, extravasation, and invasion through tissue matrix. Notably, forced overexpression of FAP promotes NK cell invasion through matrix in both transwell and tumor spheroid assays, ultimately increasing tumor cell lysis. Additionally, FAP overexpression enhances NK cells invasion into a human tumor in immunodeficient mice. These findings demonstrate the necessity of FAP in NK cell migration and present a new approach to modulate NK cell trafficking and enhance cell-based therapy in solid tumors.

**Graphical Abstract:** 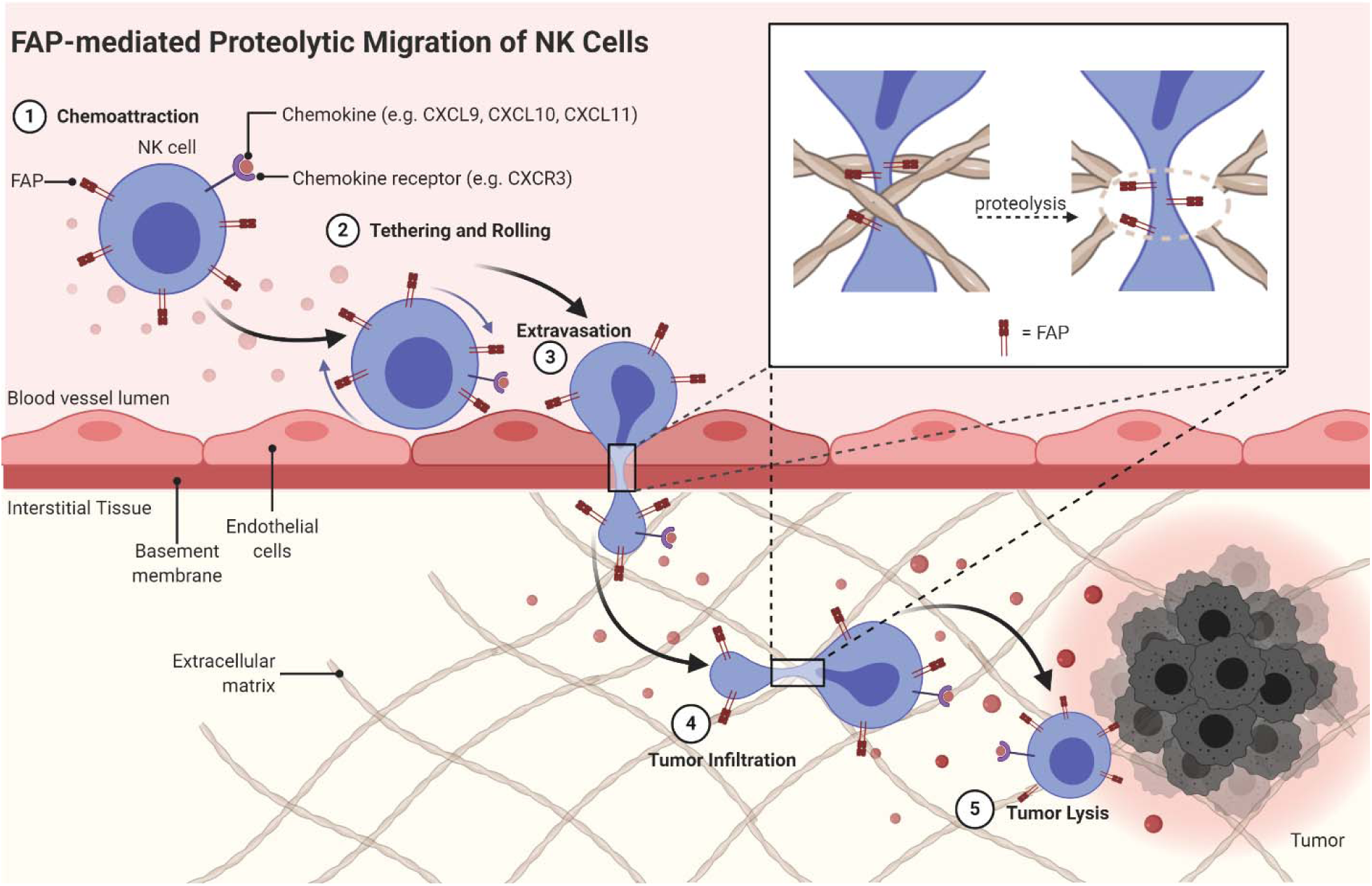

## Introduction

Natural killer (NK) cells are innate lymphoid cells that influence many physiologic and pathologic conditions through their effector and regulatory functions^1^. NK cells are canonically known to recognize and kill aberrant cells, such as virus–infected or malignant cells, using a complex detection system comprised of multiple inhibitory and activating receptors^1^. Beyond their roles as effector cells, NK cells regulate the functions of other cell types, including dendritic cells, T cells, B cells and endothelial cells, through the release of immunomodulating cytokines^2–6^. Due to their central role in the immune system and disease etiologies, efforts to manipulate NK cell activity have long been sought and developed to improve patient outcomes across many medical fields.

In cancer, patients with high tumoral NK cell content and activation have improved survival^7,8^ and response to immunotherapy^9–11^. Thus, NK cells are emerging as major targets to promote cancer immunotherapy^12^. Current NK-focused immunotherapy approaches include autologous or allogeneic NK cell transfer^13^, chimeric antigen receptor-engineered (CAR) NK cells^14^, NK cell immune checkpoint inhibitors^15^, bi- or tri-specific killer engagers (BiKEs and TriKES)^16^, and cytokine super-agonists^17^. An impediment to these therapies is inadequate NK cell homing to and/or infiltration into solid tumors.

Strategies that increase NK cell infiltration into tumors represent plausible ways to enhance NK cell-related antitumor immunotherapies. Such work has focused almost entirely on modulating NK chemokine receptors and chemoattractants^18,19^. However, lymphocyte migration depends on more than just chemotaxis. For NK cells to successfully infiltrate any tissue, including solid tumors, they must traverse complex microenvironments (e.g., extravasation from blood vessels and navigation through dense extracellular matrices)^20^. Beyond the chemokine/chemoattractant system, little is known about the mechanisms NK cells employ to physically migrate through these tissues.

Here we describe for the first time that human NK cells express fibroblast activation protein (FAP). FAP is a transmembrane serine protease primarily expressed on activated fibroblasts during wound healing or pathological conditions such as fibrosis, arthritis, and cancer^21^. FAP is primarily known for its extracellular matrix (ECM) remodeling capabilities due to its collagenase activity. Since FAP is overexpressed in diseased tissue, yet mostly absent from healthy tissue^21^, it is a promising therapeutic target in conditions like cardiac fibrosis^22^ and cancer^23^.

After identifying FAP expression by human NK cells, we used computational approaches to elucidate the function of FAP in NK cells and validated these computational findings *in vitro* using 2D assays. We then explored the impact of genetic manipulation and pharmacologic inhibition of FAP on NK cell migratory properties, extravasation, and tumor infiltration. We found that pharmacologic inhibition or deletion of FAP restrict NK cell migration, extravasation, and invasion through matrix. Conversely, forced overexpression of FAP significantly promotes NK cell invasion through matrix in both transwell and tumor spheroid assays, ultimately enhancing tumor cell lysis. Additionally, FAP-overexpressing cells showed a significantly enhanced ability to infiltrate tumors *in vivo.* These findings demonstrate the necessity of proteolytic migration in NK cell function and provide an entirely new way to enhance the anti-tumor activity of NK cells. The elucidation of FAP’s role in enhancing NK cell migration and tumor infiltration presents a promising avenue for the development of novel immunotherapeutic strategies in cancer treatment, potentially improving the efficacy of NK-cell based therapies.

## Results

### Human natural killer cells express fibroblast activation protein (FAP)

NK cells were not previously known to produce FAP; however, we detected FAP expression at the protein level in NK92 cells and three additional human NK cell lines: NKL, YT and KHYG-1 (Fig. 1A and Fig. S1 C and D). To exclude the possibility that FAP expression was specific to NK cell malignancies, we assessed FAP expression in NK cells isolated from PBMCs of five different healthy human donors and found robust FAP expression in all donor NK cells (Fig. 1B and Fig. S1E). To determine if additional human immune cell types express FAP, we assessed multiple different human B, T and monocyte cell lines for FAP expression by Western blot and found heterogeneous protein expression (Fig. 1C). This cell-line specific FAP protein expression was consistent with FAP mRNA expression as determined by analysis of RNAseq data derived from the cancer cell line encyclopedia^24^ (Fig. S1F). While we saw heterogeneous expression of FAP in B, T and monocyte cell lines, we did not detect FAP expression in healthy donor PBMC-derived B cells (CD19^+^), T cells (CD3^+^), and macrophages (CD14^+^) (Fig. 1D and Fig. S1G). Thus, FAP expression in non-NK cell lines is likely driven by their malignant biology, since FAP can be upregulated during the process of malignant transformation^21^. To further support our Western blot data, we confirmed FAP protein expression was detected on NK92 as well as normal healthy donor NK cells by immunofluorescence (Fig. 1F).

**Fig. 1.**
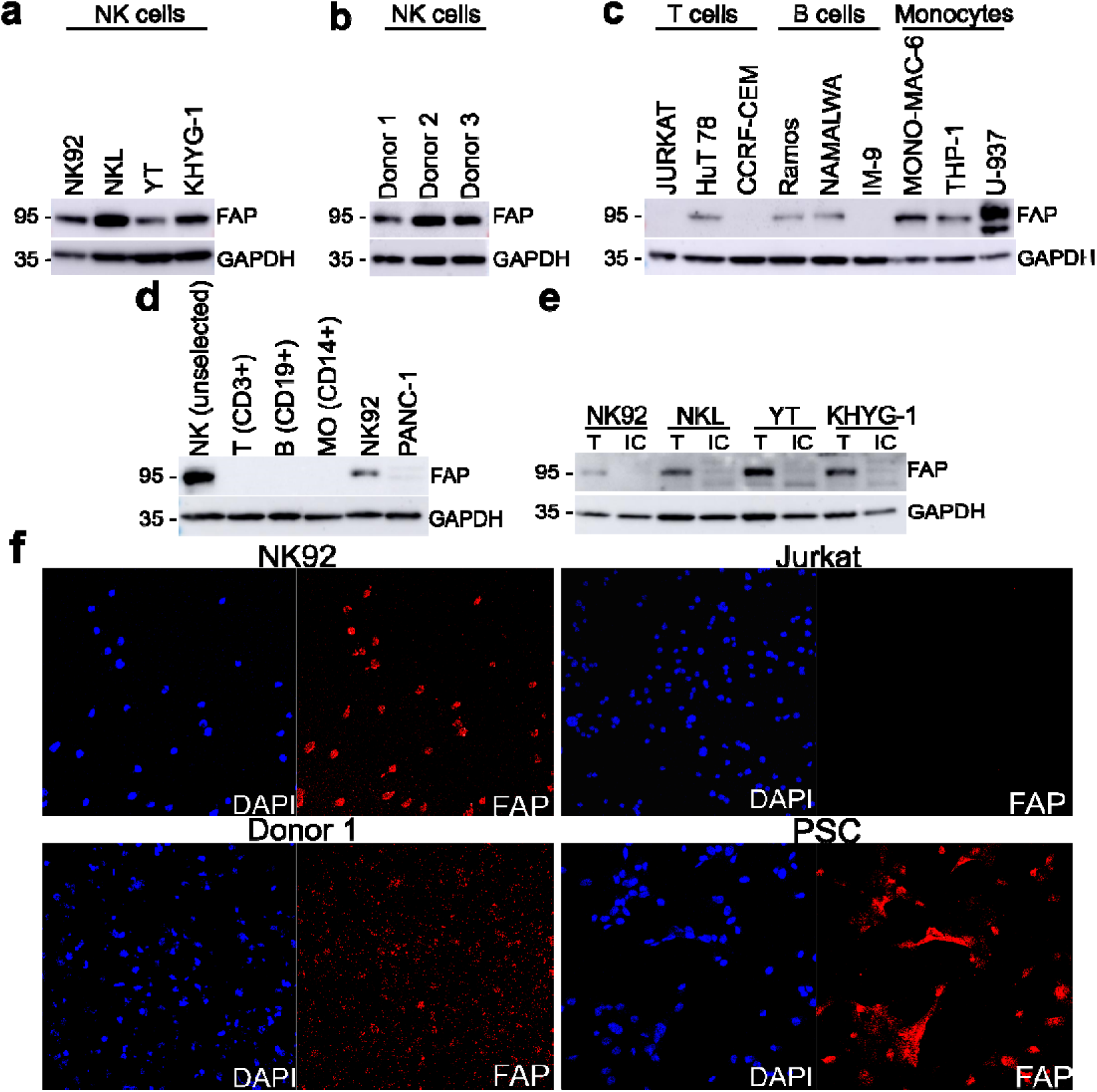
Human natural killer cells express fibroblast activation protein. (**a**) Western blot showing four distinct human NK cell lines express FAP. (**b**) Western blot showing primary NK cells isolated from PBMCs from three different healthy human donors express FAP. (**c**) Western blot showing heterogenous FAP expression in multiple human immune cell lines. (**d**) Western blot showing FAP is only expressed in donor human NK cells and not in donor human T (CD3^+^), B (CD19^+^) or monocyte (CD14^+^) cells isolated from PBMC. NK92 cell line included as a positive control and PANC-1 cell line included as a negative control. Representative of results with two different donors. (**e**) Western blot of total protein (T) and intracellular (IC) protein isolated from human NK cell lines using cell surface protein biotinylation for exclusion of surface proteins. (**f**) Immunofluorescence images showing FAP expression in NK92 and human donor NK cells. Pancreatic stellate cells (PSCs) were included as positive controls as well as Jurkat cells as negative controls. Images are representative of three separate experiments. *P* value was calculated using unpaired two-tailed t-test. \*\*\**P*<0.001, \*\*\*\**P*<0.0001. All bar plots represent mean ± SD.

Canonically, FAP is surface-expressed, so we examined FAP expression on the NK cell surface. In order to assess this, we biotinylated cell surface proteins, and then excluded them from the cell lysate via magnetic separation. We then determined that FAP is present in total cell lysate but absent from the intracellular protein lysate (Fig. 1E), demonstrating that FAP is indeed expressed on the NK cell surface.

### In NK cells, FAP gene expression correlates with extracellular matrix and migration regulating genes

We leveraged transcriptional analysis to further determine FAP’s function in human natural killer cells. In 2011, Iqbal et al. performed a gene expression array on multiple NK cell lymphoma samples and NK cell lines^25^. Using these data, we assessed FAP expression in 22 NK cell lymphomas and 11 NK cell lines (Fig. 2A) and performed a correlation analysis to assess the genes that were most positively and negatively correlated with FAP expression (Fig. 2B). The top 19 genes that were most positively correlated with FAP expression are shown in Figure 2C. We then performed GO enrichment analysis of these genes and determined that the pathways most positively correlated with FAP expression were related to ECM remodeling and cellular migration (Fig. 2D). This is consistent with the current understanding of FAP function, which is to cleave ECM components such as collagen and enhance cellular migration/invasion^21^. It is also interesting that matrix metalloproteases (MMPs) were among the top 19 genes positively correlated with FAP expression. MMPs regulate rat, mouse and human NK cell migration into collagen or Matrigel *in vitro*^26–28^. These data suggest that FAP may also regulate NK cell migration.

**Fig. 2.**
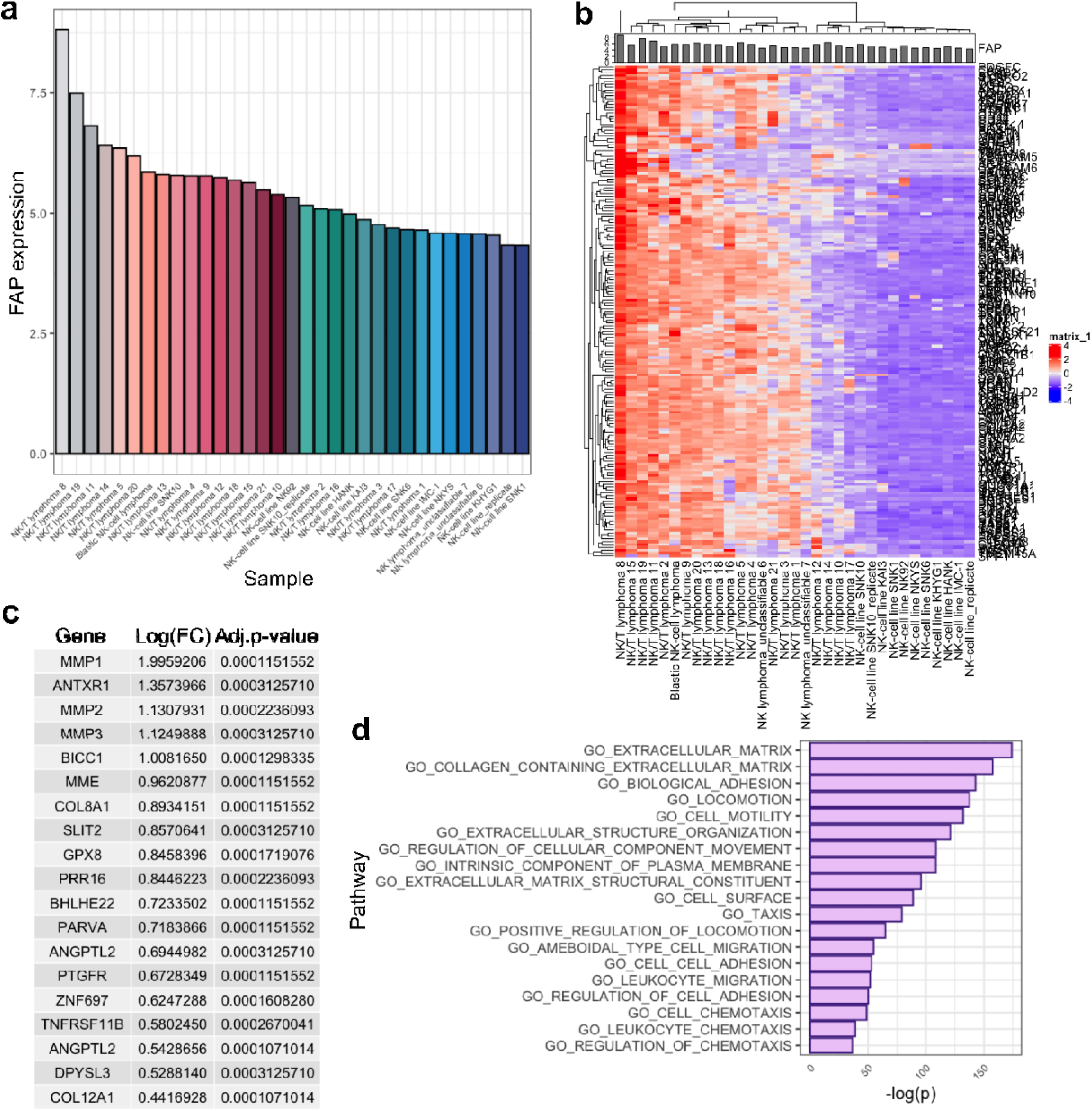
In NK cells, FAP gene expression correlates with extracellular matrix and migration-regulating genes. (**a**) Level of FAP expression in NK cell lymphomas (n=22) and NK cell lines (n=11) as determined by Affymetrix gene expression array. (**b**) Heatmap of gene expression array data. Data are shown as z-score scaled values. (**c**) Top 19 genes that are significantly correlated with FAP expression. (**d**) Top GO pathways that significantly correlate with FAP expression.

### Manipulation of FAP regulates human NK cell migration on matrix

Based on the computational analysis, we hypothesized that FAP was expressed by human NK cells to enhance migration. To test this hypothesis, we initially compared primary NK cell migration *ex vivo* in the presence and absence of an FAP-specific inhibitor (Cpd60)^29^ that inhibited FAP but not FAP’s most closely related protein, DPPIV (Fig. 3A) or the other members of the prolyl oligopeptidase family S9^29^. Cpd60 had no effect on NK cell viability (Fig. S2A).

**Fig. 3.**
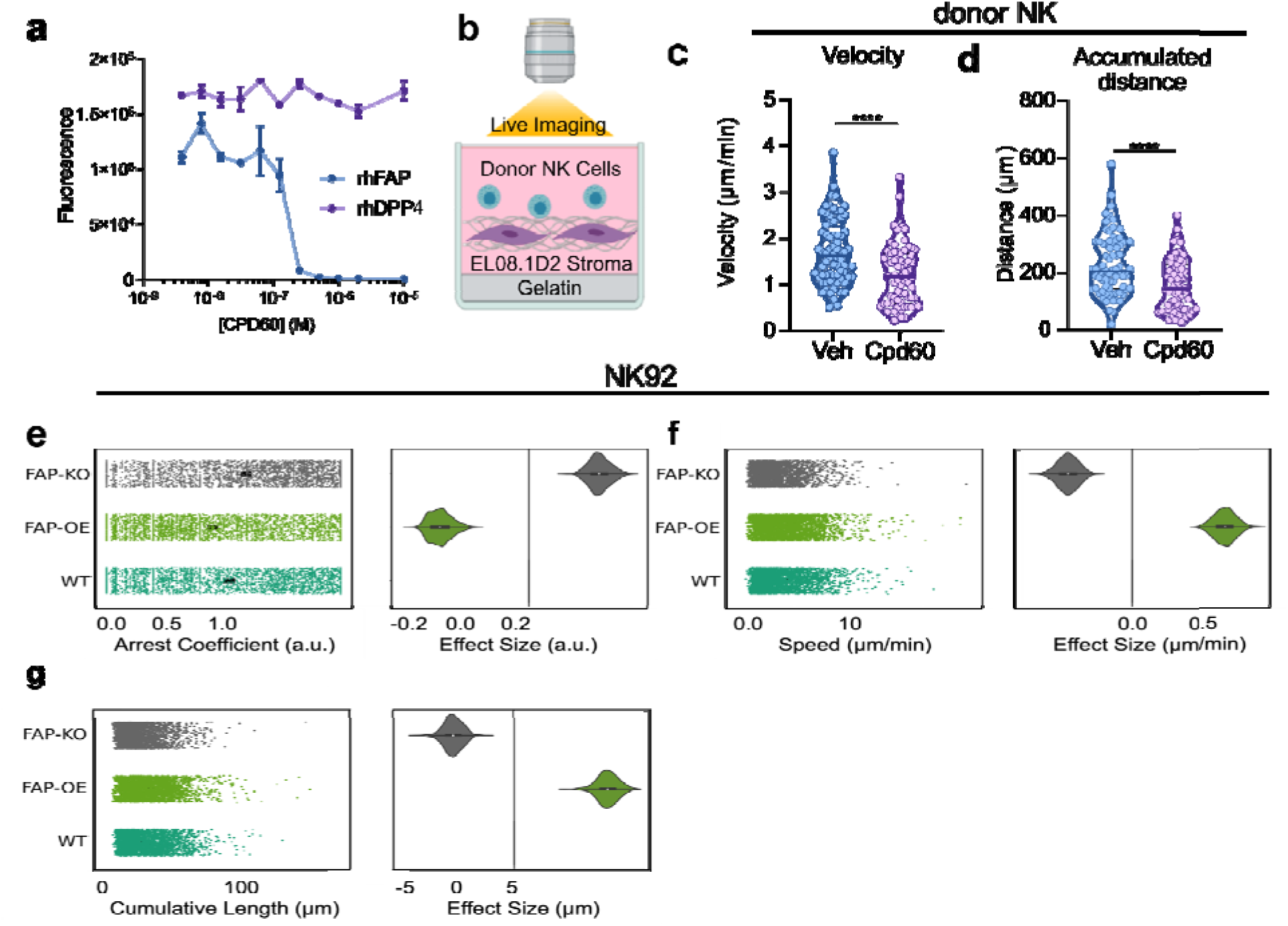
Manipulation of FAP regulates human NK cell migration on matrix (**a**) Fluorescent peptide dipeptidyl peptidase activity assay demonstrating FAP inhibitor (Cpd60) inhibits FAP but not DPPIV. (**b**) Schematic of live imaging of primary human NK cell migration on stromal cells. (**c**) The average velocity and (**d**) accumulated distance traveled by primary NK cells treated with either vehicle or 10µM Cpd60. Each point represents a single NK cell. Each condition contains 90 NK cells with 30 NK cells from three separate donors. Violin plot horizontal line represents the mean. (**e**) Arrest coefficient, defined as the frequency of time cells were found in arrest, (**f**) mean cell speed, and (**g**) cumulative track length of FAP KO NK92, FAP OE NK92, and NK92 cells. Data are shown as Plots of Difference with the wildtype condition as the reference. n=800-1800 cells per condition from 4 technical replicates.

We then cocultured primary NK cells with EL08.1D2 cells, which have previously been shown to support spontaneous NK cell migration ^30,31^ and produce ECM^32^, and live imaged them for 24 h capturing photos every 2 minutes (Fig. 3B). From these time-lapse videos we manually tracked NK cell migratory paths (Movie S1 and S2). These experiments were repeated with NK cells from three different donors, with similar results. We found that FAP inhibition with Cpd60 significantly reduced NK cell velocity (Fig. 3E) and the accumulated distance traveled by NK cells (Fig. 3F).

These findings were confirmed using an FAP knockout NK92 cell line. FAP was knocked out (FAP KO) in NK92 cells via a CRISPR-Cas9 system. This knockout was confirmed by Western blot and rt-qPCR (Fig. S3A and B). Similar to primary cells, NK92 cells were incubated on a confluent layer of EL08.1D2 stromal cells and imaged at 2 min intervals for 1-3 hours. Instead of manual tracking, cells were segmented and tracked using automated detection and tracking as described in Methods. Because of this, we were able to get data from 800-1800 cells per condition. We found that FAP KO cells displayed significantly longer arrest coefficients, defined as the frequency of time cells were found in arrest, (Fig. 3E), slower speed (Fig 3F), and lower accumulated distance (Fig 3G).

We hypothesized that if FAP KO reduced NK cell migration then FAP overexpression would increase migration. We generated a FAP-overexpressing NK92 cells line (FAP OE) using retroviral transfection. These cells were selected via the GFP expression conferred by the plasmid (Fig. S4A). This upregulation was confirmed by Western blot and RT-qPCR (Fig. S4B and C). As hypothesized, FAP OE NK92 cells displayed significantly shorter arrest coefficients (Fig. 3E), faster speed (Fig. 3F), and longer accumulated distance (Fig. 3G).

### FAP manipulation regulates the invasion of human NK cells through matrix

We then examined the impact of FAP on NK cell invasion through matrix. To analyze this, NK92 cells (NK92, FAP KO, FAP OE) were plated in the top well of a transwell chamber. CXCL9, a known NK cell chemoattractant^33^, was placed in the lower well to stimulate NK cell invasion. NK cells were allowed to invade through the membrane coated with a Matrigel barrier for 24 hours (Fig. 4A). Notably, FAP OE in NK92 cells resulted in an almost three-fold increase in invasion through Matrigel. (Fig. 4C). Additionally, knockout as well as inhibition of FAP enzymatic activity with Cpd60, an FAP specific inhibitor, resulted in a significant decrease in invasion through the Matrigel (Fig. 4B). No additional decrease was seen following treatment of FAP KO cells with Cpd60, suggesting that the decrease seen in response to Cpd60 is due to the inhibition of FAP (Fig. S5).

**Fig. 4.**
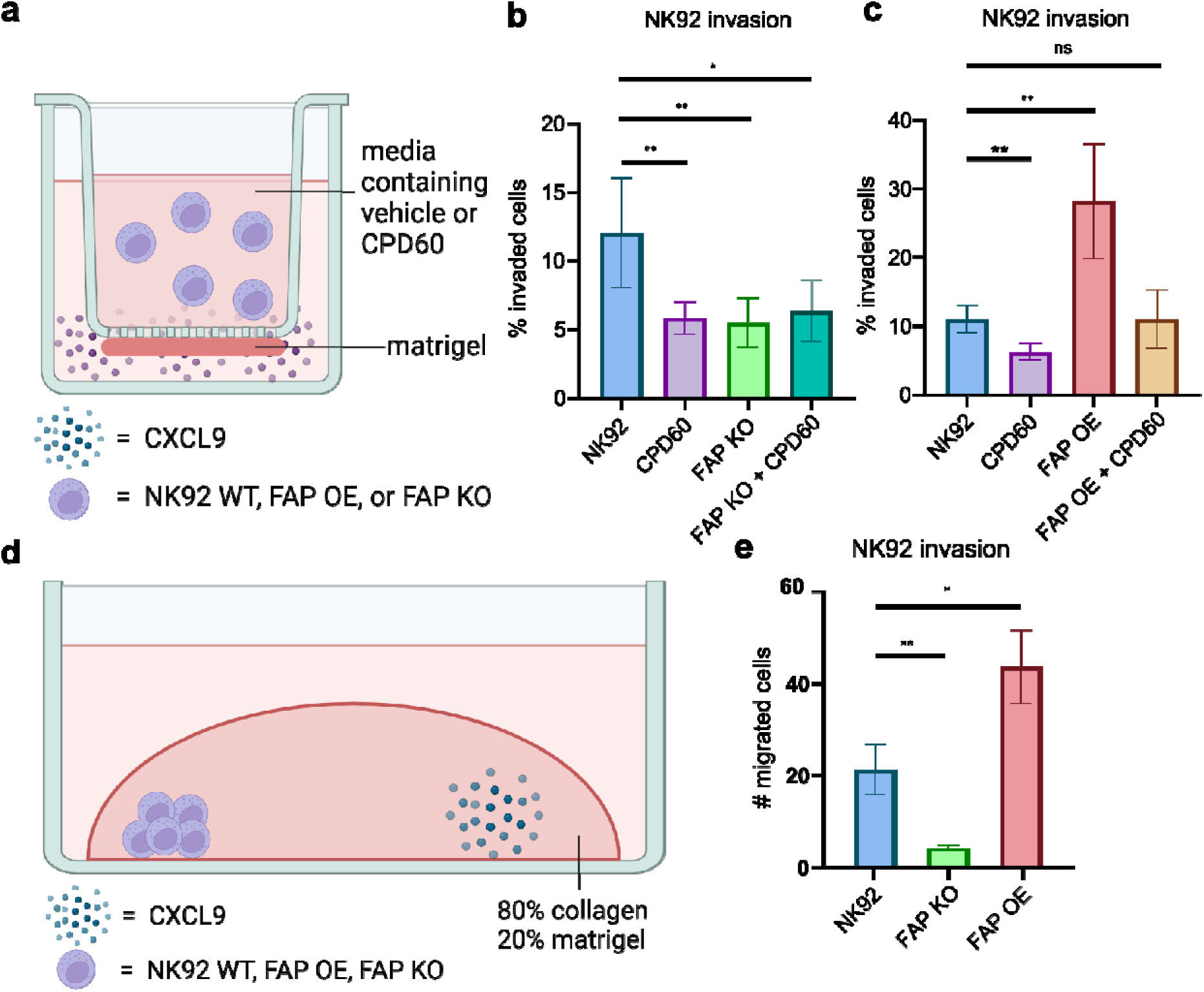
FAP manipulation regulates invasion of human NK cells through matrix. (**a**) Schematic representation of transwell assay. (**b**) Quantification of the percent of NK92 and FAP KO NK92 cells with and without Cpd60 that have invaded to the lower portion of transwell chambers. Data are from three different experiments. (**c**) Quantification of the percent of NK92 and FAP OE NK92 cells with and without Cpd60 that have invaded to the lower portion of transwell chambers. Data are from three separate experiments (**d**) Schematic representation of droplet assay. (**e**) Quantification of NK92, FAP KO NK92 and FAP OE NK92 cells migrated beyond the initial droplet. Data are from three separate experiments. *p<0.05, **p<0.01, ***p<0.001 as determined by unpaired two-tailed t-test. All bar plots represent mean ± SD.

To verify these findings, we performed a droplet assay. 2,000 NK cells (NK92, FAP KO, and FAP OE) were plated on one side of a four well LabTek plate with CXCL9 plated on the other side of the well. They were then covered in ECM and allowed to invade for 24 hours (Fig. 4D). FAP KO in NK92 cells significantly reduced invasion through matrix while FAP OE in NK92 cells significantly increased invasion through matrix (Fig. 4E).

### FAP inhibition reduces NK cell extravasation in vivo

We next set out to determine if FAP altered NK cell migratory behaviors *in vivo*. Since we could not detect FAP expression in murine NK cells (Fig. S1H) we opted to use zebrafish—a novel *in vivo* model that allows us to monitor human NK cell migratory behaviors in real-time. We injected NK92-GFP cells into the precardiac sinus of *Tg(kdrl:mCherry-CAAX)y171* zebrafish embryos that express endothelial membrane-targeted mCherry (Fig. 5A). Immediately after injection, NK cells migrated via the circulation to the caudal hematopoietic tissue (Fig. 5B) gradually disseminating throughout the rest of the zebrafish vasculature. Using confocal live-imaging, with images taken approximately every 3 minutes, we captured an NK cell crawling along the inside of the blood vessel, searching for an appropriately sized pore just prior to extravasation (Fig. 5C and Movie S3, Fig. S6). After confirming that human NK cells could migrate throughout and extravasate from zebrafish vasculature, we tested the effects of FAP inhibition on NK cell extravasation. Since fluorescent microscopy is amenable to imaging multiple fish simultaneously, we used fluorescent microscopy to quantify the effects of the FAP inhibitor Cpd60 on NK cell extravasation. We confirmed that the fluorescent microscope was capable of detecting NK cell extravasation (Fig. 5D), and then imaged 20 fish injected with NK92-GFP cells, 10 of which were bathed in 10 µM of Cpd60, and 10 fish that were bathed in vehicle. We found that FAP inhibition significantly reduced NK cell extravasation from the blood vessels (Fig. 5 E and F).

**Fig. 5.**
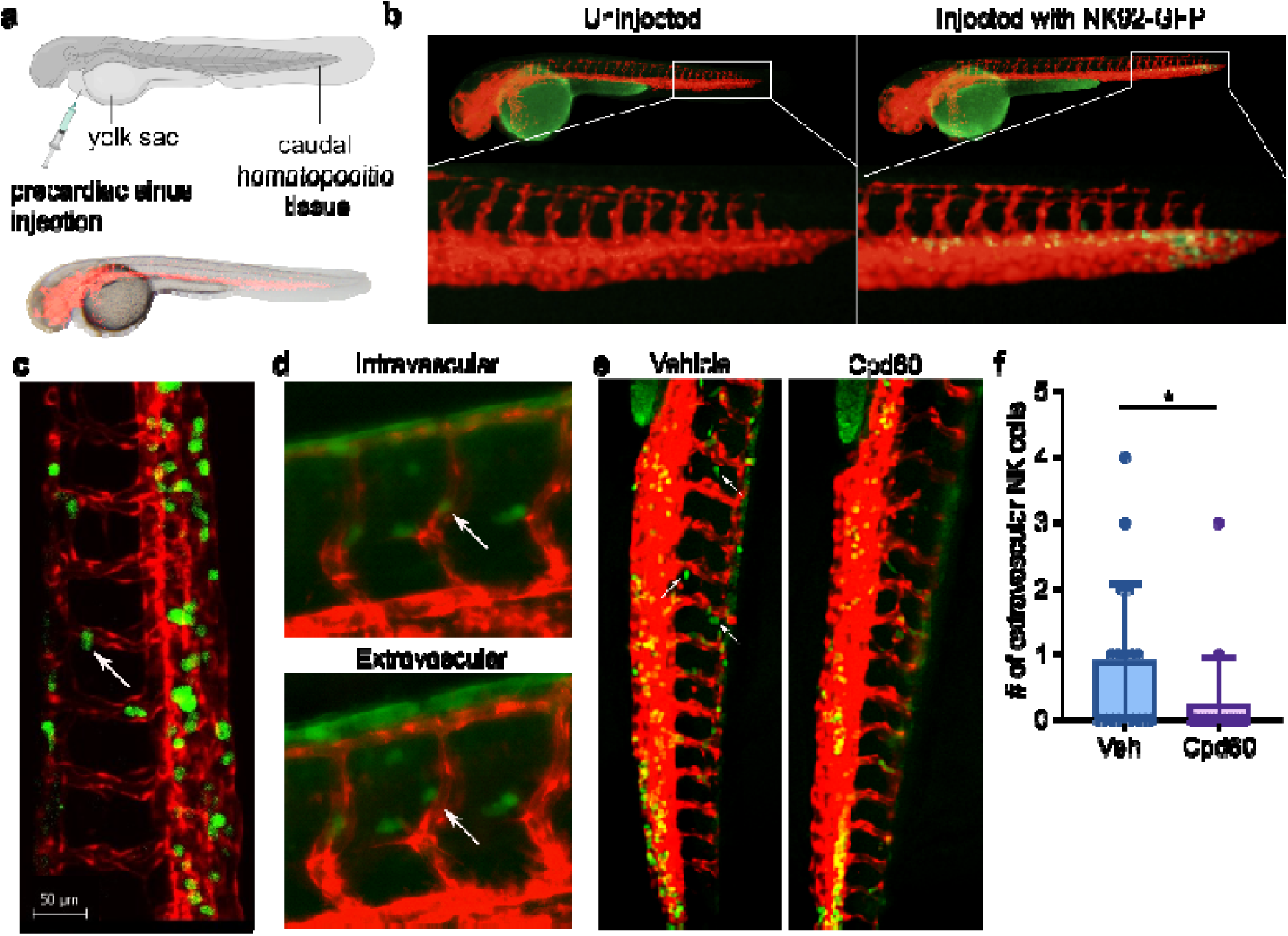
FAP inhibition reduces NK cell extravasation from zebrafish blood vessels. (**a**) Schematic representation (top) of zebrafish injections. Fluorescent and brightfield overlay image of *Tg(kdrl:mCherry-CAAX)y171* zebrafish embryos expressing endothelial membrane targeted mCherry (bottom). (**b**) Representative images of caudal hematopoietic tissue immediately after NK92-GFP injection into the precardiac sinus. (**c**) Still image taken from confocal time-lapse video demonstrating NK92-GFP extravasation from mCherry labeled vasculature. (**d**) Representative fluorescent microcopy images demonstrating NK92-GFP extravasation. Extravascular image was taken approximately 5 minutes after the intravascular image. Images were taken at 20X. (**e**) Representative fluorescent microscopy images of NK92-GFP injected zebrafish in 10 µM FAP inhibitor (Cpd60) or vehicle showing NK92-GFP cell intravascular or extravascular localization 1 hour after injection. Images were taken at 10X. (**f**) Quantification of extravascular NK92- GFP cells in zebrafish injected with NK92-GFP cells 1 hour prior to imaging. *p<0.05 analyzed by unpaired two-tailed t-test. Data are aggregated from two independent experiments, each with 10 fish per treatment condition and quantification was done blinded to treatment conditions. Bar plot represents mean ± SD.

### FAP manipulation regulates NK cell infiltration and lysis of PANC-1 cell clusters embedded in matrix

NK cells regulate tumor growth and viability, yet relatively little is known about the mechanisms NK cells employ to invade through dense tumor-related extracellular matrices. To determine if FAP activity affects NK cell infiltration into tumors, we assessed the effect of FAP inhibition on NK cell infiltration into PANC-1 clusters embedded in matrix. To accomplish this, we plated 1,000 PANC-1 cells in low-adhesion U-bottom plates and allowed them to form clusters for 24 hours. We then embedded the clusters in matrix that consisted of 80% collagen/20% Matrigel and NK92-GFP cells, and added either 10 µM Cpd60 or vehicle to the media. We live imaged the cocultures for 24 hours, capturing images every 30 minutes. Then we fixed the slides and stained for GFP by immunofluorescence to quantify the amount of NK cell infiltration into the clusters (Fig. 6A). FAP inhibition had no effect on cluster size (Fig. S7A). FAP inhibition significantly reduced NK92-GFP cell infiltration into PANC-1 clusters embedded in matrix (Fig. 6B, Movies S4-7). These experiments were repeated using PSCs with similar results (Fig. S8).

**Fig. 6.**
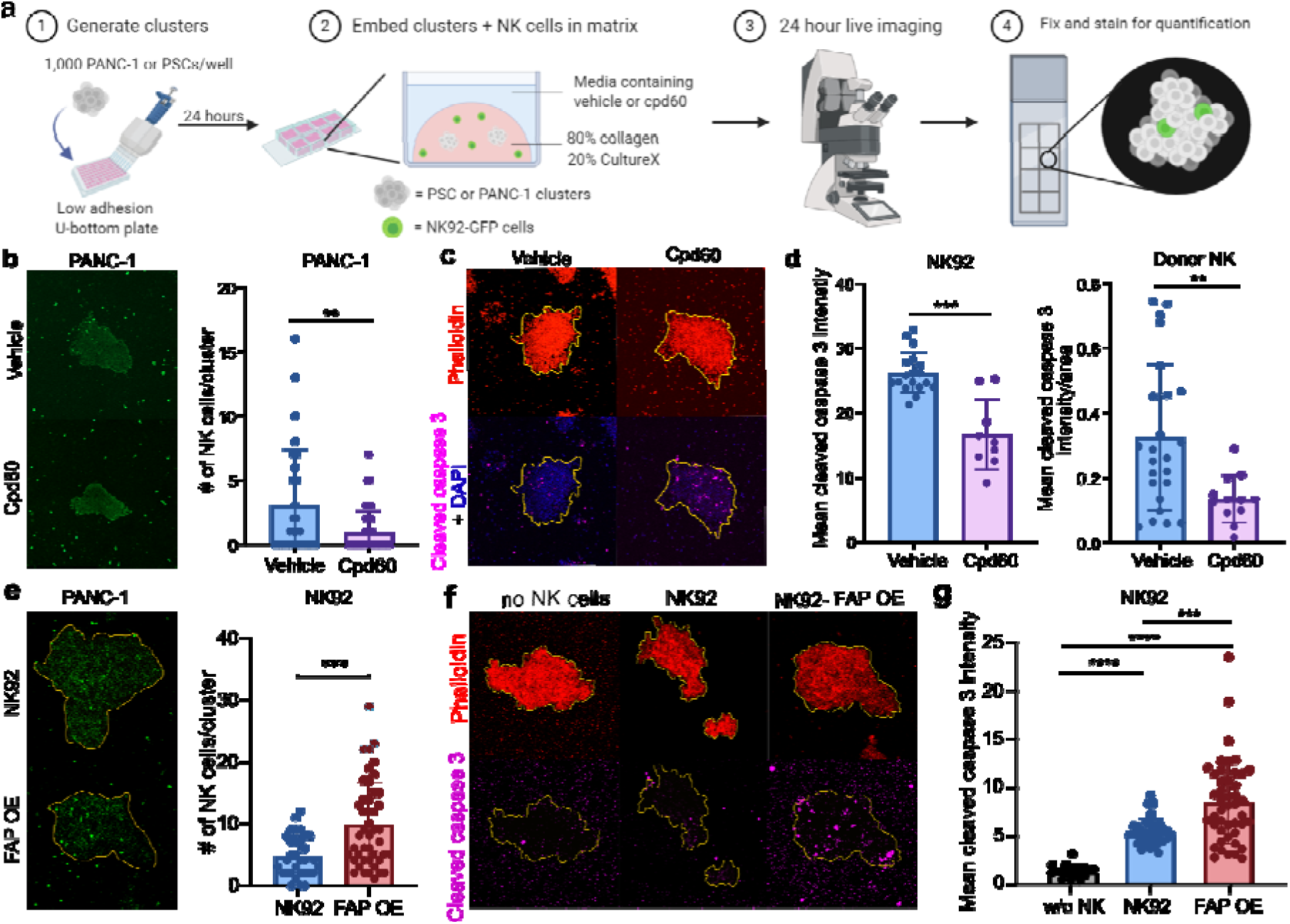
FAP manipulation regulates NK cell infiltration and lysis of PANC-1 cell clusters embedded in 3D cell matrix. (**a**) Schematic representation of experimental design. (**b**) Representative immunofluorescence images and quantification of NK92-GFP cell infiltration into PANC-1 after 24-hour coculture with vehicle or 10 µM Cpd60. PANC- 1+vehicle n = 29; PANC-1+Cpd60 n=45. PANC-1 data aggregated from two independent experiments. (**c**) Representative immunofluorescence images of phalloidin and cleaved caspase-3 staining in PANC-1 cell clusters cocultured with NK92 and vehicle or 10 µM Cpd60. (**d**) Quantification of cleaved caspase-3 intensity staining in PANC-1 cell clusters cocultured with NK92 cells or donor NK cells. Three outliers were removed from the vehicle group and one outlier was removed from the Cpd60 group. Outliers were determined by Rout method where Q = 1%. PANC-1+NK92+vehicle n = 18; PANC-1+NK92+Cpd60 n = 9; PANC-1+Donor NK+vehicle n=25, PANC-1+Donor NK+Cpd60 n =12. Donor NK cell data is aggregated data from two independent experiments that used different donors. (**e**) Representative immunofluorescence images and quantification of GFP positive NK92 cells (NK92 or FAP OE) infiltration into PANC-1 after 24-hour. PANC-1+NK92 n = 32; PANC-1+FAP OE NK92 n = 45. (**f**) Representative immunofluorescence images of phalloidin and cleaved caspase-3 staining in PANC-1 cell clusters cocultured without NK cells, with NK92, or FAP OE NK92 cells. (**g**) Quantification of cleaved caspase-3 intensity staining in PANC-1 cell clusters cocultured without NK cells, with NK92 cells, or FAP OE NK92 cells. PANC-1 without NK cells n=18; PANC-1+NK92 n = 32; PANC-1+FAP OE NK92 n = 45. Data are aggregated from two independent experiments. *p<0.05, **p<0.01, ***p<0.001 as determined by unpaired two-tailed t-test. All bar plots represent mean ± SD.

To determine if the reduced NK cell infiltration was accompanied by reduced tumor cell lysis, we repeated the PANC-1 and NK92 coculture experiment and stained the cells for actin using phalloidin and cleaved caspase-3 to identify apoptotic cells. Using the phalloidin stain we outlined the PANC-1 cell cluster, and then transposed the outline onto the cleaved caspase-3 images and quantified the intensity of cleaved caspase-3 within PANC-1 cell clusters (Fig. 6D and F). We found that FAP inhibition significantly reduced the amount of PANC-1 cell apoptosis (Fig. 6E) in 3D cultures, despite having no effect on PANC-1 cell apoptosis in 2D cell cocultures (Fig. S7B). To determine if FAP inhibition also reduced donor NK cell migration and tumor lysis, we repeated these experiments with NK cells from two donors. Since the range of PANC-1 cluster areas in the donor NK cell experiment was much wider than the range in the NK92 experiment (10-208 versus 12-70) we normalized the intensities in the donor NK cell experiment to the area of the cluster. In agreement with the NK92 cell experiments, FAP inhibition reduced donor NK cell lysis of PANC-1 cells in 3D (Fig. 6D) but not 2D (Fig. S7B). This demonstrates that FAP inhibition does not alter target cell lysis through direct impacts on NK cell cytotoxicity but rather via modulation of NK cell migration through matrix.

To determine whether FAP overexpression enhances NK cell invasion into these tumor spheroids, we repeated these experiments with NK92 cells and FAP OE NK92 cells. FAP OE NK92 cells showed significantly increased invasion into tumor spheroids and significantly increased cleaved caspase-3 expression (Fig. 6G). No increase in cytotoxicity in FAP OE NK92 cells was seen in 2D systems, suggesting the increase in apoptosis is due to an increase in NK cell invasion (Fig. S9). This suggests that FAP overexpression could be a method to enhance tumor infiltration by NK cells.

### FAP overexpression enhances NK cell infiltration into tumors in vivo

As a proof of concept experiment, we set out to determine if NK cells engineered to overexpress FAP displayed enhanced infiltration into in a human tumor murine model. To test this, we injected NK92, FAP KO NK92, and FAP OE NK92 cells intravenously into mice bearing PANC-1 pancreatic tumors. Specifically, we injected 1×10^6^ PANC-1 cells subcutaneously into NSG mice then waited until tumors were at least 100 mm^3^ in size before injecting 1×10^7^ NK92, FAP OE NK92, or FAP KO NK92 cells into the tail vein (Fig. 7A). The mice were euthanized after 24 hours and tumors were collected, fixed and stained with an anti-CD56 antibody. Tumors from mice injected with FAP OE NK92 cells had more than three times as many NK cells when compared to tumors from mice injected with NK92 cells (Fig. 7B,C).

**Fig. 7.**
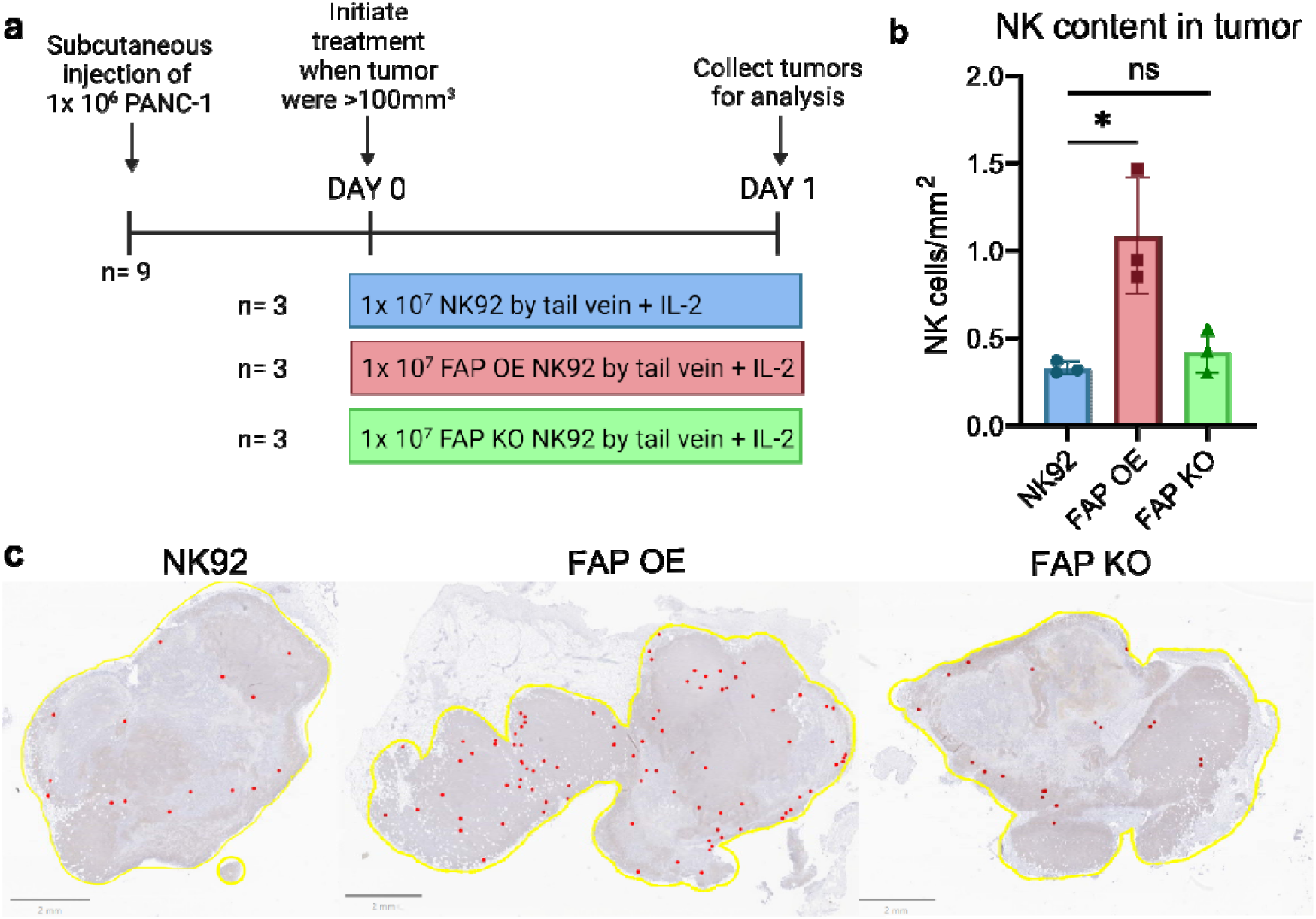
FAP overexpression enhances the infiltration of intravenously administered NK92 cells into tumors. (**a**) Schematic representation of experimental design. (**b**) Quantification of NK92, FAP OE NK92, and FAP KO NK92 cells per mm^2^ in PANC-1 tumors. (**c**) Representative hematoxylin and eosin stained images of PANC-1 tumors injected with NK92, FAP OE NK92, and FAP KO NK92 cells. Yellow indicates the border of the tumor and individual NK cells are indicated in red. n=3 mice. *p<0.05 as determined by unpaired two-tailed t-test. All bar plots represent mean ± SD.

Interestingly, there was no difference in invasion between the NK92 and FAP KO NK92 cells. Potentially because an effect size is too small to detect with our limited sample size. Alternatively, other known drivers of NK cell migration, such as MMPs, could potentially compensate for the loss of FAP^26,27,28^.

To assess NK cell trafficking to non-tumor locations, we collected and stained spleen, liver, and lung samples from each of the treated mice. There was no significant difference in NK content in any of the examined organs (Fig. S10). This suggests that under these experimental conditions NK92 cells preferentially invade into tumors rather than other organs; further supporting the notion that this technology could be implemented therapeutically.

## Discussion

Here we show that human NK cells express FAP, which is a key regulator of NK cell migration, invasion, extravasation and tumor infiltration. This novel finding significantly broadens the existing understanding of NK cell migration and tissue infiltration, and illustrates a mechanism for NK cell extravasation from blood vessels. Our findings reveal that both knockout and inhibition of FAP restrict NK cell tumor infiltration, and attenuate NK cell-mediated tumor cell lysis, underscoring the critical role of FAP-mediated migratory mechanisms in the anti-cancer activity of NK cells. Importantly, FAP overexpression enhances NK cell invasion through matrix, promoting tumor infiltration both *in vitro* and *in vivo*. Therefore, this work reveals novel insights into FAP biology and NK cell biology, with important implications for emerging NK cell-focused therapeutic strategies.

For extravasation or tissue invasion, cells must penetrate the basement membrane and interstitial tissue. During this process they are confronted by a 3D ECM that provides a substrate for adhesion and traction, as well as biomechanical resistance. For cells to traffic effectively through the ECM, which can offer narrow or non-existent pores for passage, leukocytes must adopt contracted shapes. Excessive cellular deformation can result in nuclear rupture that causes genomic damage, long-term genomic alterations and limited cellular survival. To circumvent nuclear damage, some cells employ proteolytic digestion to widen pores in the ECM^20^. Although proteolytic migration is considered less common in leukocytes versus other cell types, it has been documented. Zebrafish neutrophils and macrophages use proteolytic digestion for basement membrane transmigration^34^. Human neutrophils secrete elastase, a serine protease, to facilitate their endothelial transmigration^35^.

Unlike other immune cell types, there are few studies investigating the physical mechanisms driving NK cell migration. Decades-old research demonstrated that mouse and rat NK cell migration through Matrigel was dependent on MMPs^27,36,37^. More recent studies have used physiologic models. Putz et al. showed that heparinase regulated mouse NK cell infiltration into murine tumors^38^. Prakash et al. showed that granzyme B released from murine cytotoxic lymphocytes, including NK cells, enhanced lymphocyte extravasation via ECM remodeling, although it did not affect interstitial migration. They confirmed that a granzyme B inhibitor reduced human donor T cell transmigration through a Matrigel coated semi-permeable membrane (i.e., Boyden chamber assay)^39^. Although these authors did not assess changes in human donor NK cell migration in response to a granzyme B inhibitor, it is reasonable to assume it would be similar to that of T cell migration, since both cell types express and release granzyme B. However, our finding that FAP is expressed in human NK cells, but not in all murine NK cells or other human immune cell types (Figure 1), suggests that some migratory mechanisms can be cell-type and species-specific. Unlike these previous studies that investigated either extravasation or tumor infiltration, we investigated both and found that NK cells use the same proteolytic migration strategy for basement membrane degradation/extravasation as well as tumor tissue infiltration. We further demonstrate that defects in proteolytic migration directly impair the ability of NK cells to infiltrate and lyse tumor cells.

FAP is a well-studied protein. Although once thought to be restricted to activated fibroblasts, FAP expression has been found in additional cell types such as epithelial tumor cells^40–42^, melanocytes^43^ and macrophages^44,45^. In non-immune cells, FAP enhances cellular invasion^43,46–49^. The role of FAP in macrophages is less clear. Arnold et al. showed that in murine tumors there is a FAP^+^ minor sub-population of immunosuppressive F4/80^hi^/CCR2^+^/CD206^+^ M2 macrophages. While this study highlighted how FAP^+^ macrophages affect tumor growth, FAP’s function in these macrophages was not described^44^. Tchou et al. identified FAP^+^CD45^+^ cells in human breast tumors by immunofluorescence. They then used flow cytometry to demonstrate that a portion of these FAP^+^CD45^+^ cells were CD11b^+^CD14^+^MHC^-^II+ tumor associated macrophages. Since the flow cytometry panel used to categorize these FAP^+^CD45^+^ cells consisted of only macrophage markers, those data do not exclude the possibility that some of the FAP^+^CD45^+^ tumor cells were NK cells. In contrast to that study, we did not identify FAP expression in human macrophages (CD14^+^ cells) (Figure 1F). However, we examined circulating cells, as opposed to cells in the tumor microenvironment. Future studies are needed to further categorize FAP expression in tumor immune cell populations, potentially using multicolor immunofluorescent staining. Additionally, more studies are needed to determine if the function of FAP in FAP^+^ tumor macrophages is the same as we have described here in NK cells.

The finding that human NK cells express FAP (Figure 1D) has several clinical implications for existing FAP-targeted therapies. For example, an anti-FAP/IL-2 fusion protein has been utilized in clinical trials though the results are not yet published (NCT02627274). The proposed mechanism of action of this drug is that it targets IL-2 to FAP expressing tumor stroma, thereby limiting on-target, off-site toxicities associated with IL-2 cytokine therapy. Our findings that FAP is expressed on the NK cell surface suggests that anti-FAP/IL-2 fusion protein may also target IL-2 directly to NK cells, enhancing NK cell activation and potentially tumor clearance.

Anti-FAP CAR therapies are also in development to treat conditions such as cardiac fibrosis^50,22^, malignant pleural mesothelioma^51^, lung adenocarcinoma^52^ and other cancers^53,54^. Our data suggest that anti-FAP CAR cells may also be useful in NK cell malignancies such as aggressive NK-cell leukemia. There are potential caveats to the clinical use of anti-FAP CAR T cells. It is plausible that an anti-FAP CAR T cell could induce NK cell lysis, resulting in NK cell leukopenia in humans, this toxicity might be missed in preclinical murine models. For cancer immunotherapy, an ideal anti-FAP CAR would be engineered to target FAP expression by fibroblasts while sparing NK cells. It should be noted that Gulati et al. performed the first-in-human trial of an anti-FAP CAR T cell therapy and demonstrated that a FAP CAR T cell therapy induced stable disease for 1 year in a patient with malignant pleural mesothelioma without any treatment-terminating toxicities^51^.

Our finding that FAP regulates NK cell tissue infiltration (Figures 6 and 7) has clinical implications as well. These results imply the potential value of NK cells engineered to overexpress FAP in enhancing tumor infiltration and cell lysis.

Existing strategies aimed at enhancing NK cell infiltration into tumors rely on manipulating chemokine/receptor pathways. For example, Wennerberg et al. demonstrated that *ex vivo* expanded NK cells express higher levels of chemokine receptor CXCR3 than unexpanded NK cells which then demonstrated increased migration towards CXCL10 expressing melanomas^18^. Another approach that has been utilized is engineering NK cells to overexpress CXCR2, a chemokine receptor. This study showed that CXCR2 overexpressing NK cells had enhanced trafficking towards and lysis of renal cell carcinoma cells *in vitro*^19^. These findings suggest that these strategies to enhance NK cell migration are feasible, however, chemokine pathway-altering strategies require not only elevated expression of the chemokine receptor on NK cells, but also secretion and maintenance of chemoattractants by the tumor. Additionally, many chemoattractants recruit multiple immune cell types, including immunosuppressive cells. For example, CXCL10 is a chemoattractant for cytotoxic T lymphocytes and NK cells, but also for regulatory T cells^56^. We postulate that the ideal migration-altering therapeutic approach would increase cytotoxic immune cell infiltration in tumor masses, without influencing or even reducing immunosuppressive immune cell content in the TME. Since overexpressing FAP enhances NK92 cell tumor infiltration and lysis *in vitro* and *in vivo* (Figures 6 and 7), we speculate that engineering NK cells to overexpress FAP, either in autologous NK cell or NK CAR-NK therapies, could increase NK cell tumor infiltration and lysis. This approach is independent of tumor-associated factors, such as chemoattractant secretion, and would not be expected to induce the infiltration or expansion of immunosuppressive cell populations into the tumor microenvironment. Since proteolytic migration is required for NK cell killing of malignant cells (Figure 6), the ability to alter protease expression or activity to enhance NK cell tumor infiltration represents a potentially promising approach to altering NK cell anti-tumor activity. Future studies are needed to explore the benefit of FAP-overexpressing NK cells in preclinical models and in clinical studies, and to determine what, if any, toxicities they induce.

In this study we have demonstrated that human NK cells express FAP and that FAP directly affects NK cell migration, extravasation and tumor infiltration. These findings further the understanding of both FAP and NK cell biology. Importantly, FAP overexpression promotes the infiltration of NK92 cells into human tumor xenografts, suggesting a role for manipulating FAP expression to promote NK cell therapeutics. Future studies will determine if these novel findings have meaningful implications for NK cell-based therapy strategies currently in development.

## Materials and Methods

### Cell pellets, lines and culture

Primary human PSCs (ScienCell, cat#3830) were maintained on plastic and passaged every 1-3 days in stellate cell medium (ScienCell, cat#5301). For all experiments, PSC passage 5-9 was used. All human NK cell lines (NK92, NKL, YT and KHYG-1) and murine NK cell lines (LNK) were kindly provided by Dr. Kerry S. Campbell (Fox Chase Cancer Center, Philadelphia, PA). The NK92-GFP expressed GFP due to nucleofection with pmaxGFP according to manufacturer’s protocol (Lonza, cat#VVCA-1001). All NK cell lines were cultured as previously described^57^, tested for mycoplasma every 3-6 months and fingerprinted annually. (NKL could not be fingerprinted because it has no published profile). PANC-1 cells were cultured in 10%FBS in DMEM. Phoenix amphotropic cells were provided by Dr. Kerry S. Campbell (Fox Chase Cancer Center, Philadelphia, PA) and were cultured in 10% FBS in RPMI. The cell pellets of cell lines tested for FAP expression by Western blot (Jurkat, HuT 78, CCRF-CEM, Ramos, Namwala, IM-9, mono-mac 6, THP-1, U-937, Swiss3T3, RAW264.7, JAWSII, P815, BW5147.3, EL4 and A-20) were obtained from the Georgetown Lombardi Comprehensive Cancer Center Tissue Culture and Biobanking Shared Resource.

### Healthy donor derived cells

Fresh healthy donor NK cells were purchased from AllCells with either CD56 positive selection or CD56 negative selection (Allcells, cat#PB012-P or PB012-N). For 2D migration experiments, NK cells were enriched from peripheral blood using RosetteSep (StemCell Technologies) from healthy adult donors. T cells, B cells and monocytes were isolated from PBMCs (AllCells) using Mojosort magnetic cell separation system from Biolegend via CD3 positivity (Biolegend, cat#480133), CD19 positivity (Biolegend, cat#480105), CD14 positivity (Biolegend, cat#480093). PBMC purity was assessed using flow cytometry: CD3-APC (Biolegend, cat#300411), CD14-BV421 (Biolegend, cat#325627), CD45-FITC (BD Bioscience, cat#347463), CD56-PE (BD Bioscience, cat#555516), CD20-PE (BD Bioscience, cat#555623). For donor NK cell lysis of PANC-1 clusters, primary donor NK cells were purchased from AllCells then expanded using irradiated K562-4-1BBL-mbIL-21 (names “CSTX002”) cells kindly provided by Dr. Dean Lee according to his protocol^58^.

### Generation of FAP Overexpressing Cells

Overexpression of FAP was induced in NK92 by retroviral transduction. Phoenix amphotropic cells were transfected with the pBMN plasmid containing the FAP gene (received from vectorbuilder) using Lipofectamine and Plus reagent (Life Technologies) as previously described^59^. Supernatants were collected from these cells after growing for 48 hours in Opti-MEM media (Life Technologies). The supernatant was mixed with Lipofectamine and Plus reagent and added to 2×10^6^ NK92 cells in a 6-well plate. These cells were centrifuged for 45 min at 2000xg. This process was repeated two consecutive times and cells were flow sorted for GFP positivity three days after the final transduction.

### Generation of NK92 FAP Knockout Cells

Knockout of FAP in NK92 cells was accomplished by CRISPR using nucleofection, as previously described^60^. 2µL of CAS9 RNP (Horizon Discovery, cat#CAS12205) and 2µL FAP sgRNA (Horizon Discovery, cat#SQ-003829-01-0002) were incubated together for 15 minutes at room temperature. The sgRNA complexes were then added to 1×10^6^ NK92 cells resuspended in 16uL of P3 nucleofection buffer (Lonza). The nucleofection mixture was transferred to a 16-well strip for nucleofection in the Lonza 4D Nucleofector using the pulse code CM-138. The 20µL nucleofection mixture was then added directly to 1mL of pre-warmed NK media. This process was repeated an additional time and cells were used 72 hours later.

### FAP Activity Assay

One day prior to assay, 5,000 PSCs/well were added to 96 well flat clear bottom white polystyrene TC-treated microplates (Corning, cat#3610). The following day, PSC media was aspirated off and 50 µL of NK92 cells (lacking GFP) were added to each well containing PSCs at a 4:1 E:T ratio and incubated overnight at 37°C. 100 mM stock of dipeptidylpeptidase substrate (Acetyl-Aka-Gly-Pro-AFC) (Anaspec, CatAS-24126) was made by resuspending lyophilized substrate in DMSO. On the day of the assay, DMSO stock was then diluted 1:1000 in FAP activity assay buffer (50 mM Tris-BCl, 1 M NaCl, 1 mg/mL BSA, pH 7.5). A standard curve was generated using rFAP (R&D systems, 3715-SE-010). 50 µL of rFAP standard ranging in concentration from 0.03125-2ug/mL was added to wells in triplicate. 50 µL of substrate was added to each well and the plate was incubated for 5 minutes at 37°C. The plate was read on a PerkinElmer EnVision Multimode Plate Reader with 390-400 nm excitation and 580-510 nm emission wavelengths. The final concentration of FAP per well was calculated using the standard curve. Data were compiled and assessed for statistical significance using GraphPad Prism 9.

### PSC-NK92 Coculture Assay

PSCs were plated one day prior to assay at 100,000 cells/well in a 6 well collagen coated plate. NK92 cells were added at 1:1 or 4:1 effector to target (E:T) ratios and cocultured for 3-4 hours. Each well contained 50% v/v NK and PSC media and 1% v/v IL-2. Following incubation, nonadherent cells were collected. Adherent cells were washed 2X with PBS and then trypsinized with 0.05% trypsin. After detachment trypsin was quenched with equal volume PSC media and cells were collected, pelleted and washed 2X with PBS then resuspended in 600 µL of 1% BSA. Cells were immediately sent for nonsterile flow sorting of GFP+ from GFP- using the BD FACS Aria Ilu cell sorter in the Georgetown Lombardi Comprehensive Cancer Center Flow Cytometry and Cell Sorting Shared Resource (FCSR).

### Annexin V NK cell lysis study

One day prior to assay, PSCs or PANC1 cells were stained with DiI. If donor NK cells were used, they were stained with DiO prior to the experiment. Cells were then plated as described for the PSC-NK92 coculture assay. Following incubation period of 4 hours, all cells from a single well were collected and washed 2X with PBS. Samples were then processed by the FCSR using the Alexa Fluor 647 Annexin V and Sytox Blue staining (Biolegend). Flow data were analyzed using FloJo (v10.4.1) and statistics was performed using GraphPad Prism 9.

### RNA Isolation and rt-qPCR

RNA was isolated using the PureLink RNA Mini Kit (Ambion, cat#12183020). The RNA concentration was measured using NanoDrop 8000 (Thermo Fisher Scientific). cDNA was generated from 20-100 ng of RNA using the GoTaq 2-step RT-qPCR System (Promega, cat# A6110). qPCR was performed with SYBR Green on a StepOnePlus real-time PCR system (Applied Biosystems). Gene expression was normalized to HPRT and analyzed using 1/ΔCt method.

Primers sequences:

*FAP*: (F: ATGAGCTTCCTCGTCCAATTCA; R: AGACCACCAGAGAGCATATTTTG)

*HPRT*: (F: GATTAGCGATGATGAACCAGGTT; R: CCTCCCATCTCCTTCATGACA)

### Western Blot

Western blots were performed as previously described^57^. Western blots were conducted using anti-FAP (ab207178, abcam) at concentrations of 1:1000 diluted in 5% milk in PBST. Secondary antibody was anti-rabbit IgG, HRP linked (Cell Signaling, cat# 7074S) at 1:1000. Antibody was validated with additional anti-FAP antibodies (MyBiosource, cat#MBS303414 and abcam, cat#ab53066). GAPDH antibody (Cell Signaling, cat#5174S) was used at 1:10,000. The secondary antibody was anti-rabbit IgG, HRP linked (Cell Signaling) used at 1:5000. Chemiluminescent substrate (Pierce, cat#32109 or cat#34094) was used for visualization.

### Immunofluoresence

5×10^5^ cells were plated on coverslips for 2 hours in a 12-well plate. Cells were fixed for 15 minutes at room temperature with 4% paraformaldehyde, washed with PBS, and permeabilized with 0.5% Triton X-100 for 15 minutes at room temperature. The cells were washed with PBS and then blocked with 1% BSA for 30 minutes. These cells were then incubated with primary antibody for 1 hour at room temperature. Immunofluorescence was conducted using anti-FAP antibody (Santa Cruz, sc-65398) at a concentration of 1:500. The cells were washed three times with PBS. They were then incubated with Alexa Fluor 647 anti-mouse secondary antibody (Thermofisher, A21236) at a concentration of 1:1000. The cells were again washed three times with PBS. They were then incubated with DAPI diluted 1:1000 for 20 minutes at room temperature. They were again washed three times and then mounted on a slide using ProLong Antifade Mountant (Invitrogen, cat#10144). Antibody was validated with additional anti-FAP antibody (ab207178, abcam). These slides were then imaged on a Leica SP8 AOBS microscope in the LCCC Microscopy and Imaging Shared Resource (MISR).

### Cell Surface Biotinylation

Cell surface biotinylation of NK92, NKL, YT and KHYG-1 cells was performed with the Pierce Cell Surface Protein Isolation kit (Thermo Scientific, cat#89881) according to the manufacturer’s protocol. In brief, 4×10^8^ cells were pelleted and washed with cold PBS then incubated with EZ-LINK Sulfo-NHS-SS-biotin for 30 min at 4°C followed by the addition of a quenching solution. Another 1×10^6^ cells were collected and saved for total cell western blotting. Cells were lysed with lysis buffer (500 μL) containing the complete protease inhibitor cocktail (Roche, cat#11697498001). The biotinylated surface proteins were excluded with NeutrAvidin agarose gel (Pierce, 39001). Samples were diluted 50 ug in ultrapure water supplemented with 50 mM DTT. Lysates were subjected to Western blotting with the anti-FAP antibody described above.

### Gene expression analyses of NK cell lines

NK lymphoma and cell line gene expression was downloaded from GEO (GEO accession GSE19067)^25^ using R version 3.6.2 and read using affy in Bioconductor^61^. Non-NK cell samples were excluded from analysis. Heatmap was created using ComplexHeatMap version 2.1.1^62^. Correlation analysis was performed using limma in Bioconductor^63^. Gene set enrichment analysis was performed using GO enrichment^64^.

### 2D cell migration studies

2D migration studies were done as previously reported^31,32,65^. In brief, EL08.1D2 stromal cells were grown to a confluent monolayer on flat-bottomed 96 well ImageLock plates (Essen Bioscience) pre-coated with 0.1% gelatin (Stemcell Technologies). For imaging primary cells, 10 µM of Cpd60 in RPMI media was added to the chamber 15 min before imaging. Freshly isolated human NK cells or 5,000 NK92 cells (NK92, FAP KO, FAP OE) were imaged in 96-well plates on the IncuCyte ZOOM Live-Cell Analysis System (Essen Bioscience) at 37°C in the phase-contrast mode (10× objective). Cells were allowed to settle for 30 min prior to beginning imaging every 2 minutes for 1-3 hours in an Incucyte Zoom using brightfield settings.

### Automated cell tracking and analysis

Exported TIFF stacks from Incucyte images were segmented using the Cyto2 trained network provided by Cellpose {Stringer, 2021 #3} using a classification object diameter of 7. btrack {Ulicna, 2021 #11} was used to track segmented cells between frames. Data was analysed using cellPLATO^66^. A HDF5 file containing segmented masks and tracks for each cell was generated for each TIFF stack and saved. Custom Python functions were used to make 29 separate shape, migration, and clustering measurements per timepoint per cell. Cell tracks were filtered by area (40-300 µm^2^) and by the number of timepoints a cell is segmented (5 to 1800).

### Manual cell tracking and analysis

Manual tracking of live cells was done using the manual tracking feature in Fiji^67^. Tracks were plotted using the Chemotaxis plugin of Fiji. Cells that were in the field of imaging for fewer than two frames were discarded, as were cells which were non-adherent or floating.

EL08.1D2 cells were used as de facto fiducial markers to ensure that neither they nor the microscope stage was drifting and causing apparent NK cell movement. Length and displacement measurements were derived directly from tracked cells and graphed using GraphPad software. Velocity data was obtained by dividing the total track length by the time of imaging.

### Transwell assay

Matrigel was diluted 1:4 in NK media. 50µL of this mixture was plated on the underside of a 5µm pore transwell insert (Corning, cat#CLS421). This was allowed to solidify for 20 minutes at room temperature. 2×10^5^ cells in 200µL media were plated in the top well of the plate. 100ng/mL CXCL9 (R&D systems, cat# 392-MG) was added to 400µL media plated in the lower well of the plate. The cells were allowed to migrate for 24 hours and the number of cells in the bottom well was counted using a hemocytometer.

### Droplet assay

Cells were stained with DiO prior to the experiment. 2,000 cells were resuspended in 1µL of ECM mixture and plated on one end of a well on a 4 well Labtek plate (Thermo Scientific, cat#154917). 0.8ng of CXCL9 (R&D systems, cat# 392-MG) was resuspended in 2µL of ECM and plated on the other end of a well. The ECM mixture consisted of 20% growth factor reduced Matrigel (Corning, 10-12 mg/ml stock concentration, #354230) and 80% rat tail collagen type I at 3mg/mL (Gibco, A1048301). The two droplets were then covered in 75µL of ECM and was allowed to solidify for 45 minutes at 37°C. 800µL of NK media was then added to the well and the cells were allowed to migrate for 24 hours. The entire slide was then imaged on the Olympus IX-71 Inverted Epifluorescent Microscope at 5X in the LCCC Microscopy Shared Resource and the number of cells that had migrated beyond the initial droplet were counted using FIJI.

### Zebrafish studies

Zebrafish studies were conducted in accordance with NIH guidelines for the care and use of laboratory animals and were approved by the Georgetown University Institutional Animal Care and Use Committee. Zebrafish husbandry, injections, and mounting was performed by the Georgetown-Lombardi Animal Shared Resource. Two day post fertilization stage *Tg(kdrl:mCherry-CAAX)* embryos were anesthetized with 0.016% tricaine (Sigma-Aldrich, St. Louis, MO, USA) in fish water (0.3g/L Sea Salt, Instant Ocean, Blacksburg, VA) and were injected with 100-200 NK92-GFP cells into the precardiac sinus using an air driven Picospritzer II microinjector (General Valve/Parker Hannifin) under a stereoscope. Following injection, embryos with cells in the caudal hematopoietic tissue were selected for analysis and mounted in 1.5% agarose plus 0.011% tricaine in fish water. Fish were maintained at 33°C until imaging.

Confocal imaging was performed on a Leica SP8 AOBS microscope in the Georgetown-Lombardi Microscopy Shared Resource. Widefield fluorescent imaging was performed on a Keyence BZ-X inverted microscope. Images were taken at 10X across multiple z-stacks. Z-stack images were compressed using full focus and haze reduction in Keyence BZ-X software. NK extravasation quantification was performed by counting the number of GFP cells outside red vasculature. NK extravasation quantification was performed blinded to the treatment conditions. Graphs of resulting data and statistical analysis was generated using Graphpad Prism 9.

### 3D cluster studies

3D clusters were generated, embedded and stained as previously described^68,69^. In brief, clusters were generated by plating 1,000 cells per well into 96-well Nunclon Sphera low adhesion plates (Thermo Scientific, cat#174925) and incubated overnight at 37°C. The following day, 6 clusters were embedded into an ECM containing 2,000 NK cells and were plated into one well of a Nunc Lab-Tek II 8-well chamber slide (ThermoScientific, cat#154534PK). The ECM mixture consisted of 20% growth factor reduced Matrigel (Corning, 10-12 mg/ml stock concentration, #354230) and 80% rat tail collagen type I at 3mg/mL (Gibco, A1048301). Cells were either imaged for the following 24 hours every 30 minutes using a Zeiss LSM800 scanning confocal microscope enclosed in a heated chamber supplemented with CO_2_ or allowed to incubate overnight at 37°C. After 24 hours, cells in matrix were fixed with 5.4% formalin for 1 hour, permeabilized with 0.5% Triton-X and blocked using goat serum. For invasion assays, NK-92-GFP cells were stained with anti-GFP (ThermoFisher, cat#A-11122). For the cell lysis assays, clusters were stained using anti-cleaved caspase-3 (Cell Signaling, cat#9661). Hoechst 33342, phalloidin, and secondary antibodies labeled with Alexa Fluor 488 nm, 546 nm, 647 nm, or 680 nm (Invitrogen) were used.

### Animal studies

10 NSG (NOD.Cg-Prkdc Il2rg /SzJ) mice were divided into 3 groups of 3 with one kept as a negative control. We inoculated animals with 1×10^6^ human PANC-1 cells by subcutaneous injection. When tumors were <100mm, three mice were injected with 1×10^7^ NK92 cells and 4,000 IU IL-2, three were inject with 1×10^7^ FAP OE NK92 cells and 4,000 IU IL-2, three were inject with 1×10^7^ FAP KO NK92 cells and 4,000 IU IL-2, and one was injected with saline by IV tail vein as a negative control for staining. 24 hours after injection, mice were euthanized and the tumor, lung, liver, and spleen were excised. The tumors and organs were submitted to the histopathology and tissue shared resource core at Georgetown. Slides were stained using an anti CD-56 antibody (Abcam, ab133345) at 1:800. Slides were then blinded and NK cells were manually counted using Qupath.

## Supplementary Materials

Fig. S1. Human NK cells express catalytically active fibroblast activation protein.

Fig. S2. Effects of Cpd60 on cell viability.

Fig. S3. FAP knockout in NK92 cells

Fig. S4. FAP overexpression in NK92 cells

Fig. S5. Effects of empty pBMN plasmid and NTC on NK92 invasion

Fig. S6. NK92-GFP cells probing for permissive sites to exit from zebrafish vasculature (red).

Fig. S7. Effects of Cpd60 on cluster area and NK cell lysis of target cells in 2D.

Fig. S8. Effects of Cpd60 on NK cell infiltration into PSC clusters.

Fig. S9. Effects of FAP manipulation on NK92 cytotoxicity

Fig. S10. NK cell infiltration into organs

Movie S1: Tracking of primary NK cell migration in vehicle using phase-contrast live imaging.

Images were taken 30 minutes apart for 24 hours. Each color track represents the migration path of a single NK cell.

Movie S2: Tracking of primary NK cell migration in 10 uM FAP inhibitor (Cpd60) using phase-contrast live imaging. Images were taken 30 minutes apart for 24 hours. Each color track represents the migration path of a single NK cell.

Movie S3: Time-lapse video of a NK92-GFP crawling along and extravasating from red zebrafish vasculature. Time-lapse images were taken 3 minutes apart for approximately 6 hours using confocal microscopy.

Movie S4: Time-lapse video of NK92-GFP cells migrating into PANC-1 clusters embedded in 3D matrix with vehicle. Images taken every 30 minutes for 24 hours using confocal microscopy.

Movie S5: Time-lapse video of NK92-GFP cells migrating into PANC-1 clusters embedded in 3D matrix with 10 uM FAP inhibitor (Cpd60). Images taken every 30 minutes for 24 hours using confocal microscopy.

Movie S6: Time-lapse video of NK92-GFP cells migrating into PSC clusters embedded in 3D matrix with vehicle. Images taken every 30 minutes for 24 hours using confocal microscopy.

Movie S7: Time-lapse video of NK92-GFP cells migrating into PSC clusters embedded in 3D matrix with 10 uM FAP inhibitor (Cpd60). Images taken every 30 minutes for 24 hours using confocal microscopy.

## Acknowledgments

We would like to thank Dr. Kerry Campbell for providing NK cell lines, the pBMN plasmid, and continuous intellectual and technical support; Dr. Dean Lee for providing the NK feeder cells; the Georgetown Lombardi Comprehensive Cancer Center Tissue Culture and Biobanking, Flow Cytometry and Cell Sorting, Genomics and Epigenomics, Histopathology and Tissue, and Microscopy and Imaging Shared Resources. Graphical abstract and schematics were created with BioRender.com

## Author contributions

A.A.F., R.E.M. and L.M.W. conceived the idea, designed the study and obtained funding. A.A.F. and R.E.M. wrote the manuscript, conducted experiments and analyzed the data. E.F.M. and E.J.F. performed the computational analysis and E.F.M generated the computational figures. A.N. and G.P. assisted in designing and performing 3D migration experiments. E.G. assisted in conducting the zebrafish studies. S.A.J. assisted with experiment design. P.V. provided FAP inhibitor Cpd60. E.M.M. performed and analyzed the 2D migration studies and assisted with study design.

## Competing interests

The authors have no competing interests to declare.

## Code availability statement

The authors declare that there is no custom code in this manuscript.

## Data availability statement

The public datasets analyzed in this paper are available at GEO accession GSE19067, doi: 10.1038/leu.2010.255. The authors declare that all other data supporting the findings of this study are available within the paper or its supplementary information files. All other relevant data are available from the corresponding author upon request.

## Funding

This work is supported by grants from the National Institute of Health National Cancer Institute F30 CA239441(A.A.F.), R01 CA50633 (L.M.W.), P30 CA51008 (L.M.W), and NIH-NIAID R01AI137073 to E.M.M

## Notes

### Competing Interest Statement

The authors have declared no competing interest.

### Summary of Updates

We have added experiments utilizing genetically engineered NK cells and murine animal models to confirm our findings.

