## Supplemental Material for "Fibroblast activation protein regulates natural killer cell migration, extravasation and tumor infiltration"

**Supplementary Materials**

Fig. S1. Human NK cells express catalytically active fibroblast activation protein.

Fig. S2. Effects of Cpd60 on cell viability.

Fig. S3. FAP overexpression in NK92 cells

Fig. S4. FAP knockout in NK92 cells

Fig. S5. Effects of empty pBMN plasmid and NTC on NK92 invasion

Fig. S6. NK92-GFP cells probing for permissive sites to exit from zebrafish vasculature (red).

Fig. S7. Effects of Cpd60 on cluster area and NK cell lysis of target cells in 2D.

Fig. S8. Effects of Cpd60 on NK cell infiltration into PSC clusters.

Fig. S9. Effects of FAP manipulation on NK92 cytotoxicity

Fig. S10. NK cell infiltration into organs

Movie S1: Tracking of primary NK cell migration in vehicle using phase-contrast live imaging. Images were taken 30 minutes apart for 24 hours. Each color track represents the migration path of a single NK cell.

Movie S2: Tracking of primary NK cell migration in 10 uM FAP inhibitor (Cpd60) using phase-contrast live imaging. Images were taken 30 minutes apart for 24 hours. Each color track represents the migration path of a single NK cell.

Movie S3: Time-lapse video of a NK92-GFP crawling along and extravasating from red zebrafish vasculature. Time-lapse images were taken 3 minutes apart for approximately 6 hours using confocal microscopy.

Movie S4: Time-lapse video of NK92-GFP cells migrating into PANC-1 clusters embedded in 3D matrix with vehicle. Images taken every 30 minutes for 24 hours using confocal microscopy.

Movie S5: Time-lapse video of NK92-GFP cells migrating into PANC-1 clusters embedded in 3D matrix with 10 uM FAP inhibitor (Cpd60). Images taken every 30 minutes for 24 hours using confocal microscopy.

Movie S6: Time-lapse video of NK92-GFP cells migrating into PSC clusters embedded in 3D matrix with vehicle. Images taken every 30 minutes for 24 hours using confocal microscopy.

Movie S7: Time-lapse video of NK92-GFP cells migrating into PSC clusters embedded in 3D matrix with 10 uM FAP inhibitor (Cpd60). Images taken every 30 minutes for 24 hours using confocal microscopy.

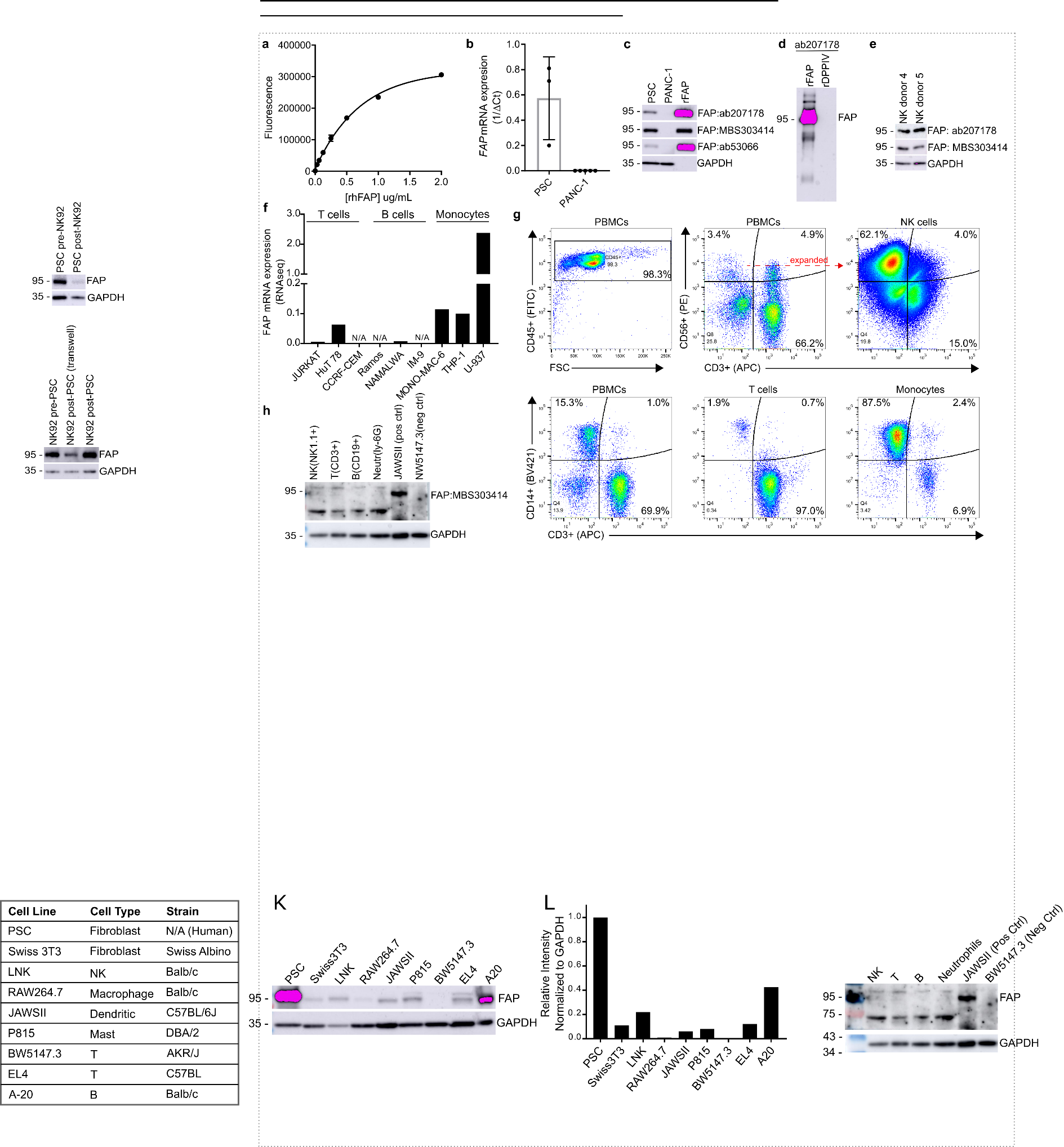
**Supplementary Fig. 1**. Human NK cells express catalytically active fibroblast activation

protein. (**a**) Standard curve for fluorescent peptide substrate assay demonstrating increased fluorescence with increasing concentration of recombinant human FAP. (**b**) Controls for qRT-PCR analysis of FAP expression in PSCs (positive control) and PANC1 cells (negative control). PSCs results determined from 3 independent experiments. PANC-1 level determined from 6 independent experiments. Data represents mean ± SD. (**c**) Controls for western blot comparing three different anti-FAP antibodies’ ability to detect FAP in PSCs (positive control), in PANC-1 (neg control) and rhFAP. (**d**) Controls for western blot showing that antibody selected for additional studies (ab2017178) detects recombinant human FAP but not FAP’s most similar protein, recombinant human DPPIV. (e) Western blot showing FAP expression in two additional primary human samples using two different anti-FAP antibodies. (f) FAP mRNA expression in multiple human leukocyte cell lines as determined by RNAseq (data derived from cancer cell line encyclopedia). N/A indicates cell lines absent from the CCLE database. (**g**) Flow cytometry analysis assessing purity of primary donor immune cells. (**h**) Western blot assessing FAP expression in flow sorted C57BL/6 splenocytes (NK, T, B, and neutrophils), in JAWSII murine cell line (positive control) and in NW5147.3 murine cell line (neg control).

**Supplementary Fig. 2**. Effects of Cpd60 on cell viability. (**a**) CellTiterBlue cell viability assay
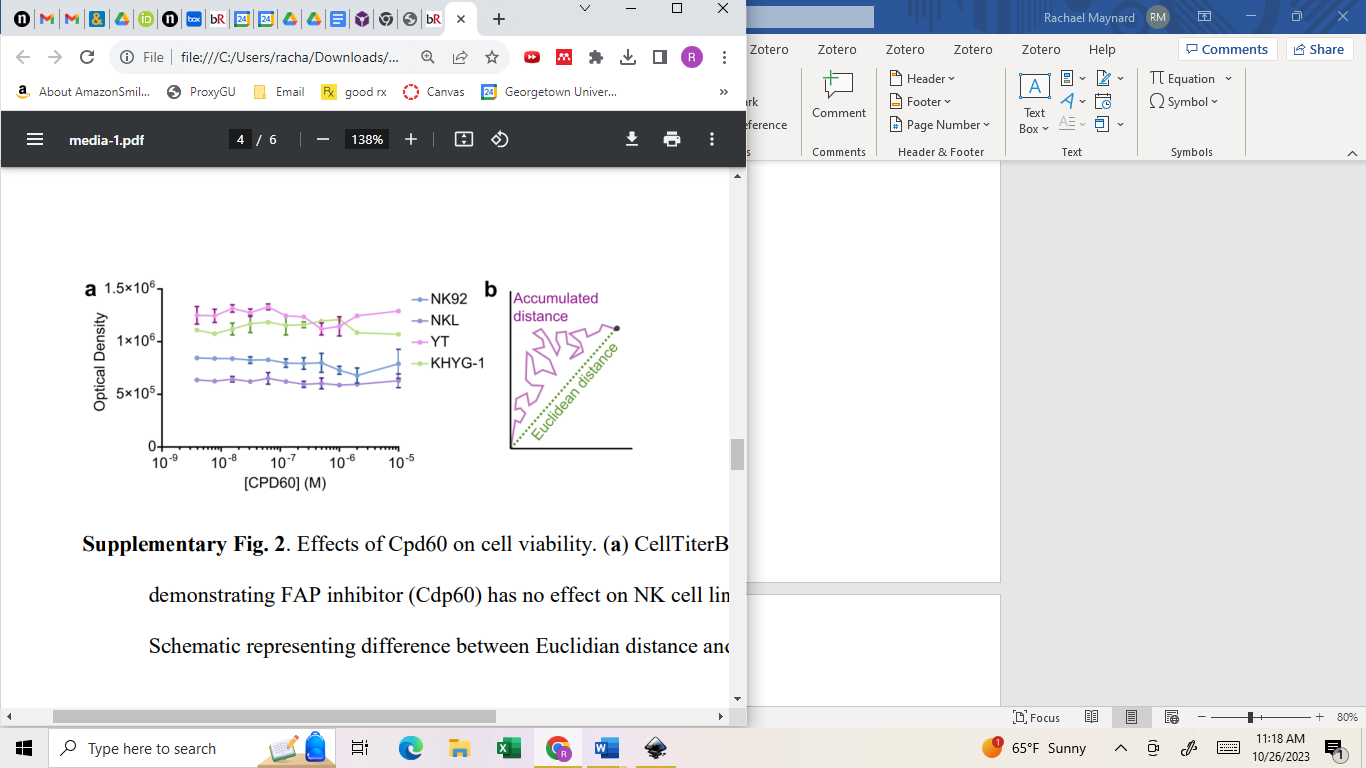

demonstrating FAP inhibitor (Cdp60) has no effect on NK cell line viability. (**b**) Schematic representing difference between Euclidian distance and accumulated distance.

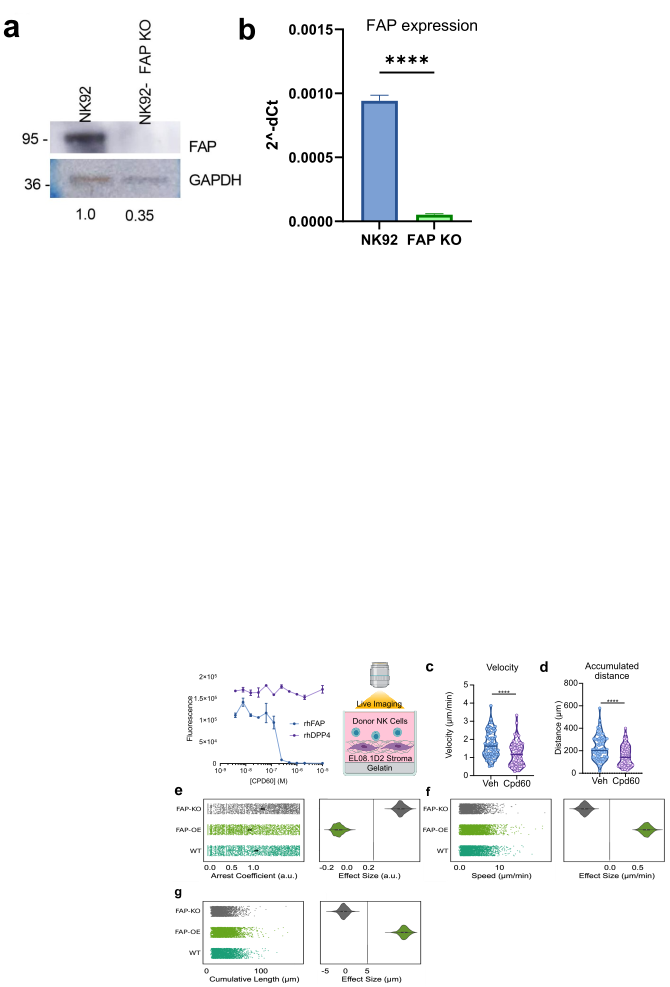

**Supplementary Fig. 3.** FAP knockout in NK92 cells. (**a)** Western blot showing FAP protein

expression in wild type NK92 and FAP KO NK92. This image is representative of three separate experiments (**b)** qPCR results showing FAP RNA expression in NK92 and FAP KO NK92. n=3. ****p<0.0001 as determined by unpaired two-tailed t-test. All bar plots represent mean ± SD.

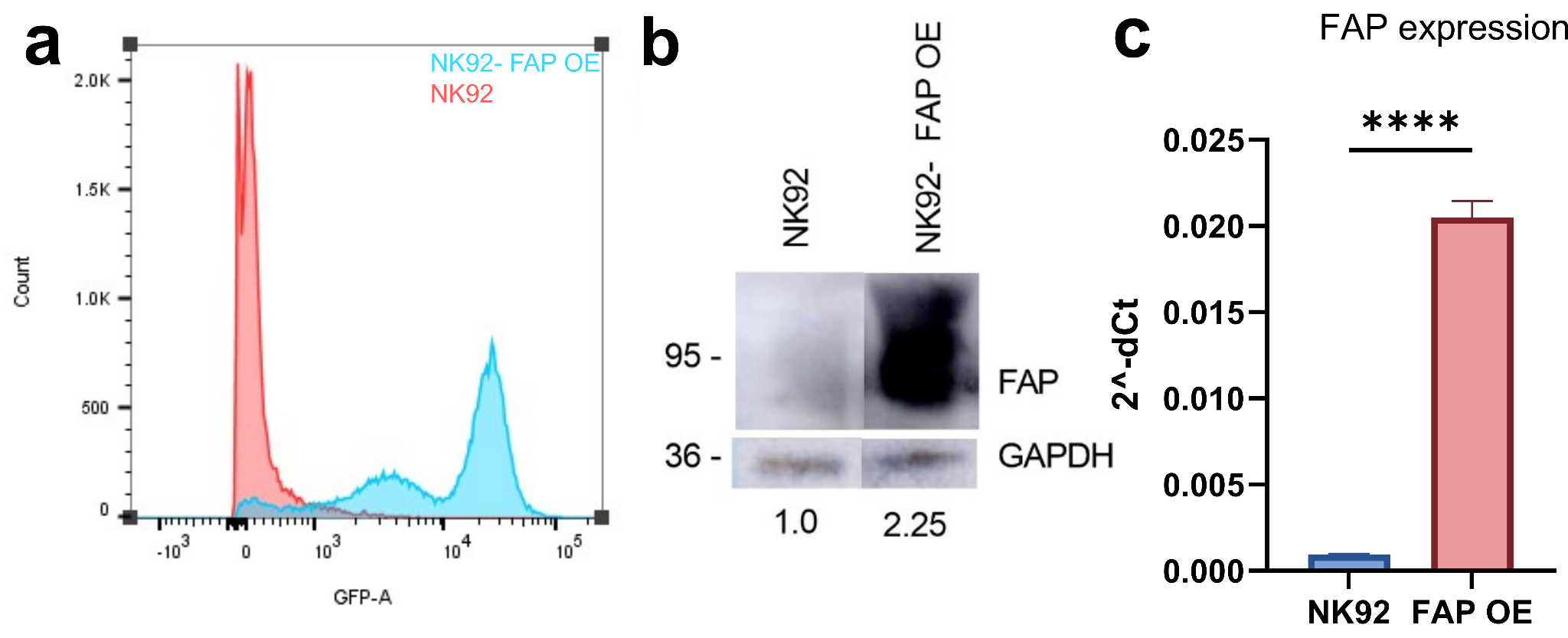
**Supplementary Fig. 4.** FAP overexpression in NK92 cells. (**a)** Flow cytometry results showing GFP expression in both NK92 and FAP OE NK92. Data are representative of three separate experiments. (**b**) Western blot showing FAP protein expression in wild type NK92 and FAP OE NK92. This image is representative of three separate experiments (**c)** qPCR results showing FAP RNA expression in NK92 and FAP OE NK92. n=3. ****p<0.0001 as determined by unpaired two-tailed t-test. All bar plots represent mean ± SD.

**Supplementary Fig. 5.** Effects of empty pBMN plasmid and non-targeting control (NTC) on
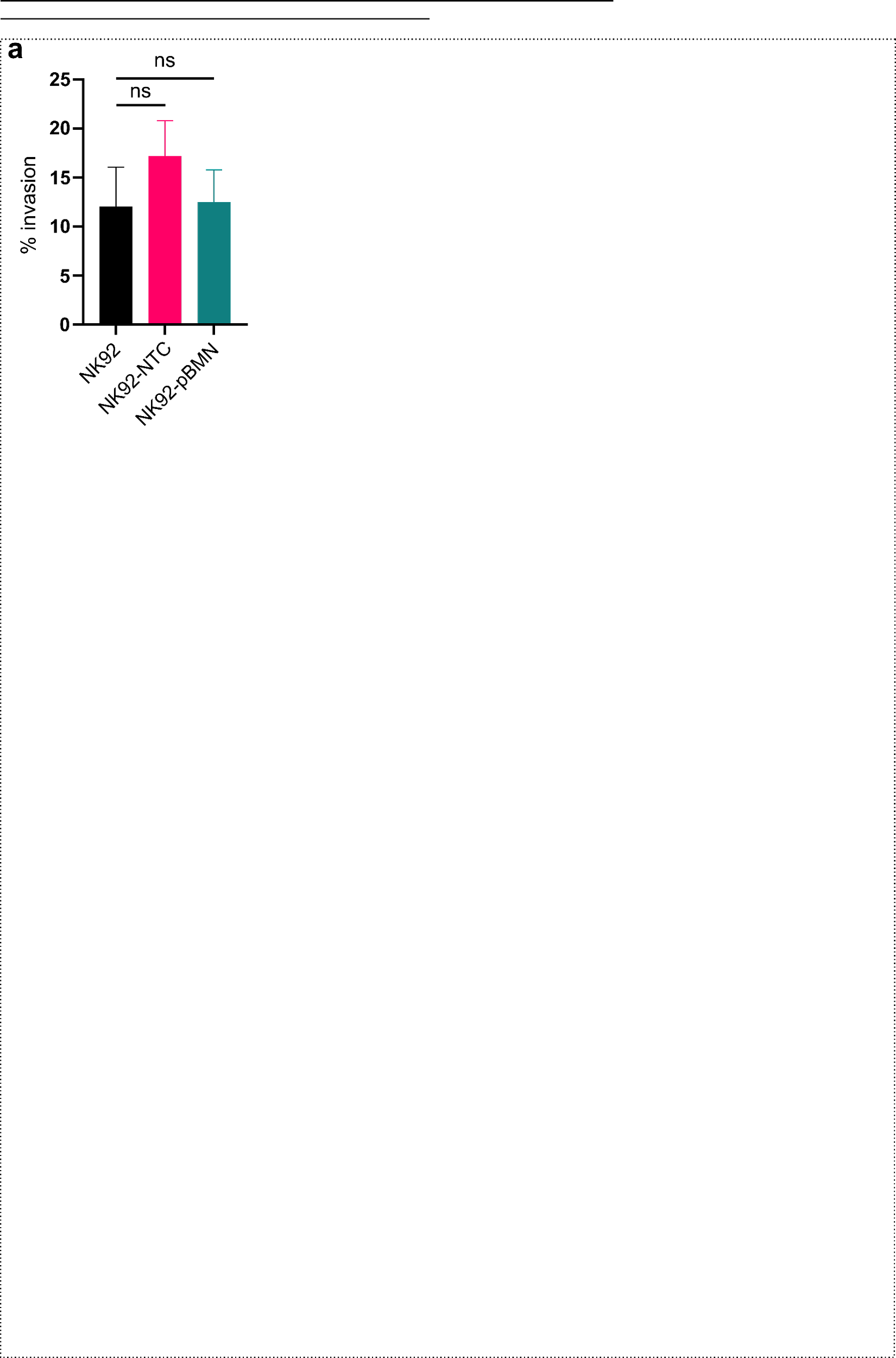

NK92 cells. (**a**) Quantification of the percent of wild type NK92, NK92 nucleofected with NTC, and NK92 cells transfected with empty pBMN plasmid that have invaded to the lower chamber of transwell. Data are from three different experiments. Ns= not signifcant as determined by unpaired two-tailed t-test. All bar plots represent mean ± SD.

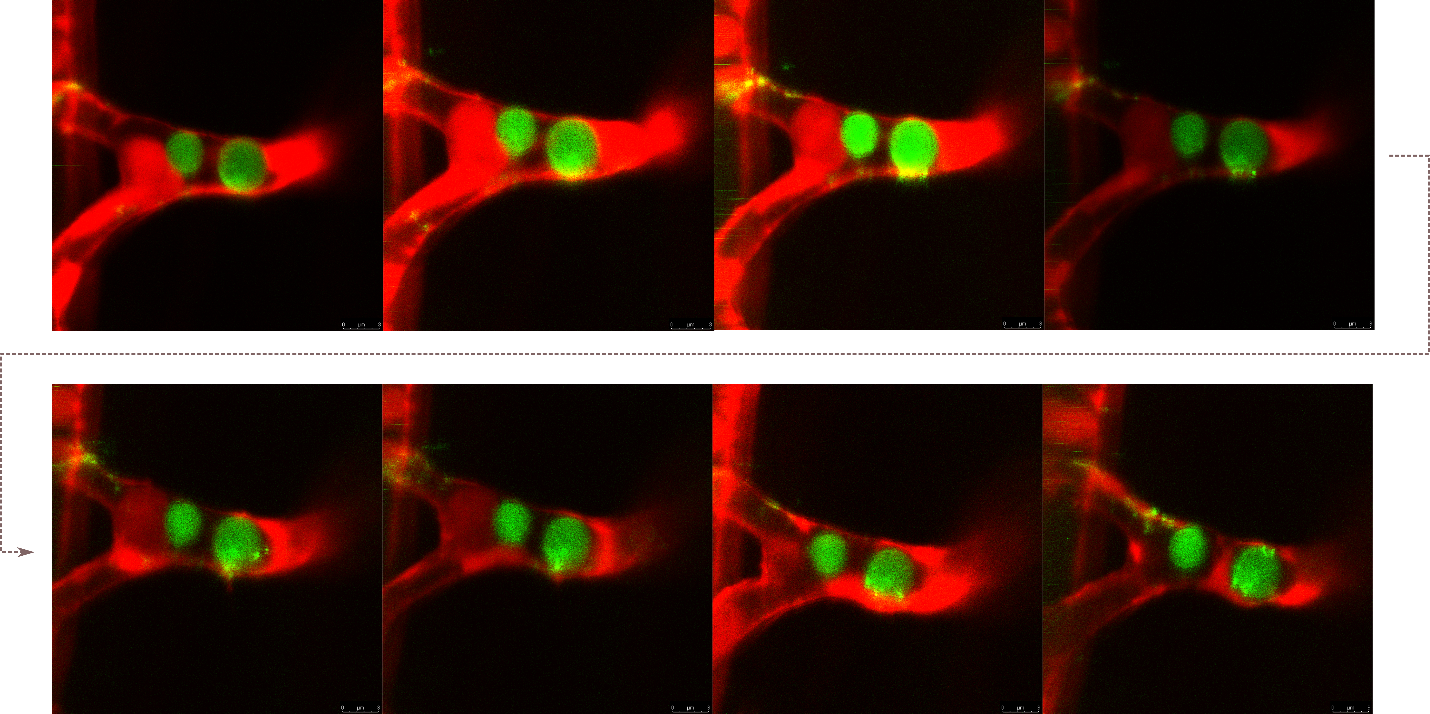
**Supplementary Fig. 6**. NK92-GFP cells probing for permissive sites to exit from zebrafish

vasculature (red).

**Supplementary Fig. 7**. Effects of Cpd60 on cluster area and NK cell lysis of target cells
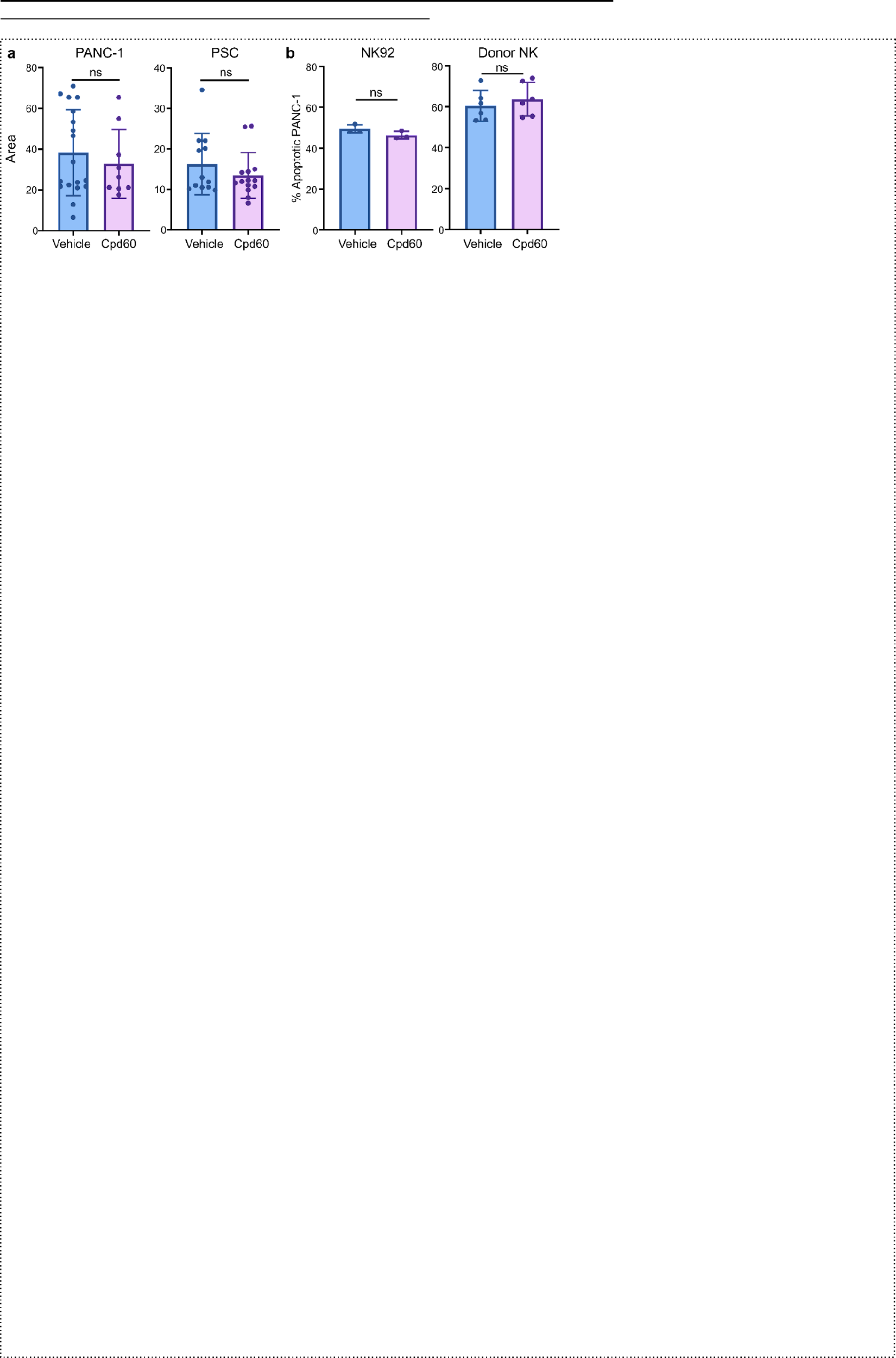

in 2D. (a) Average area of PANC-1 or PSC clusters embedded in 3D matrix with vehicle or 10uM FAP inhibitor (Cpd60). ns=not significant as determined by two-tailed unpaired t-test. PANC-1+vehicle n = 18; PANC-1+Cpd60 n = 9; PSC+vehicle n = 12, PSC+Cpd60

n = 14. (b) Impact of 10 uM Cpd60 on NK92 and donor NK cell lysis of PANC-1 cells in

2D culture as determined by annexin flow cytometry assay. The culture ratio of NK to

PANC-1 was 4 to 1. Cells were cocultured for 4 hours. NK92+vehicle n = 3;

NK92+Cpd60 n =3; donor NK+vehicle n= 6, donor NK+Cpd60 n= 6. Donor NK cell

experiment is data combined from two separate donors. All plots represent mean ± SD. two-tailed t-test. All bar plots represent mean ± SD.

**Supplementary Fig. 8.** Effects of Cpd60 on NK cell infiltration into PSC clusters. (**a**)
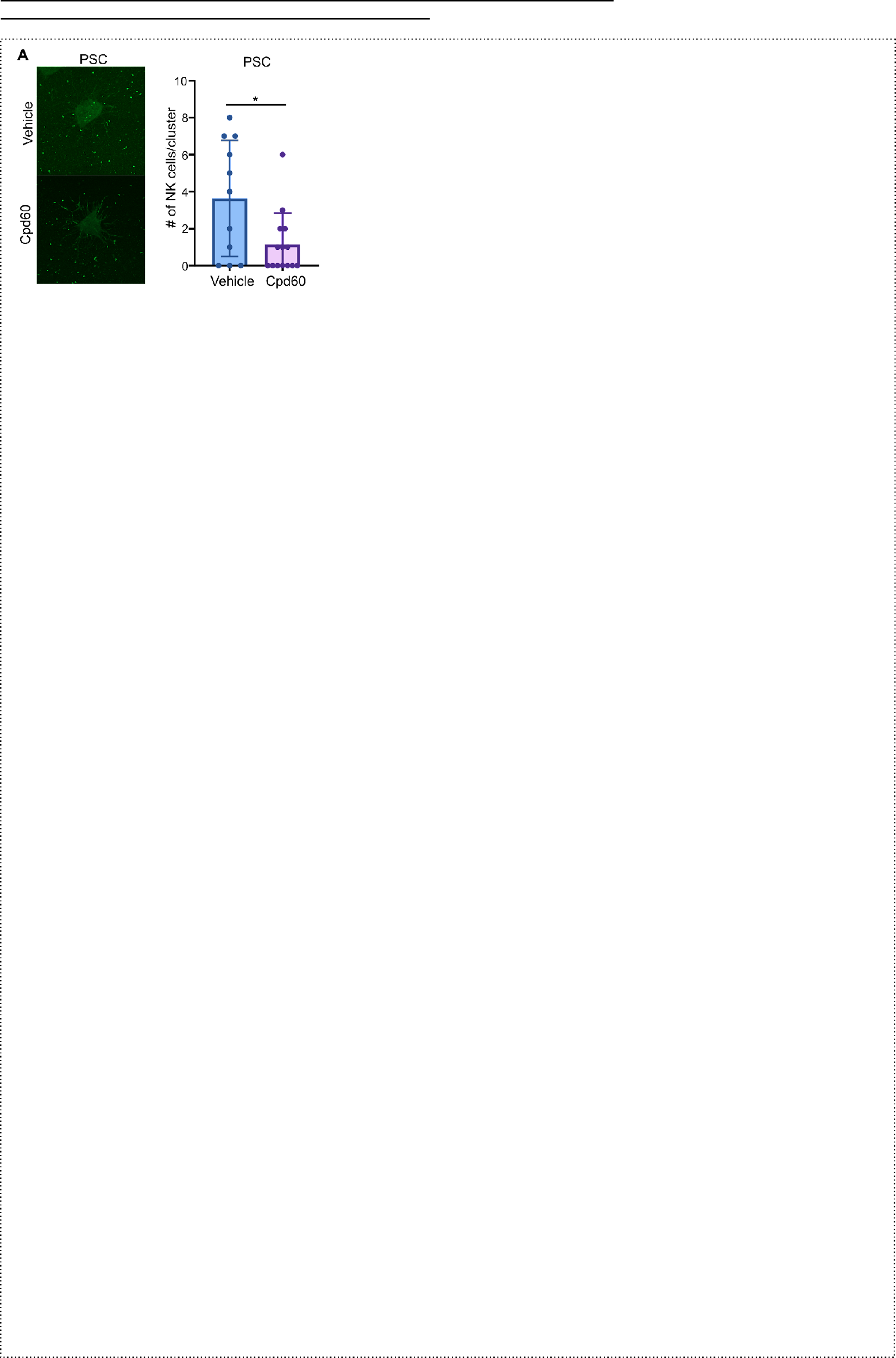

Representative immunofluorescence images and quantification of NK92-GFP cell infiltration into PSC clusters after 24-hour coculture with vehicle or 10 uM Cpd60. PSC+vehicle n = 11; PSC+Cpd60 n=14.

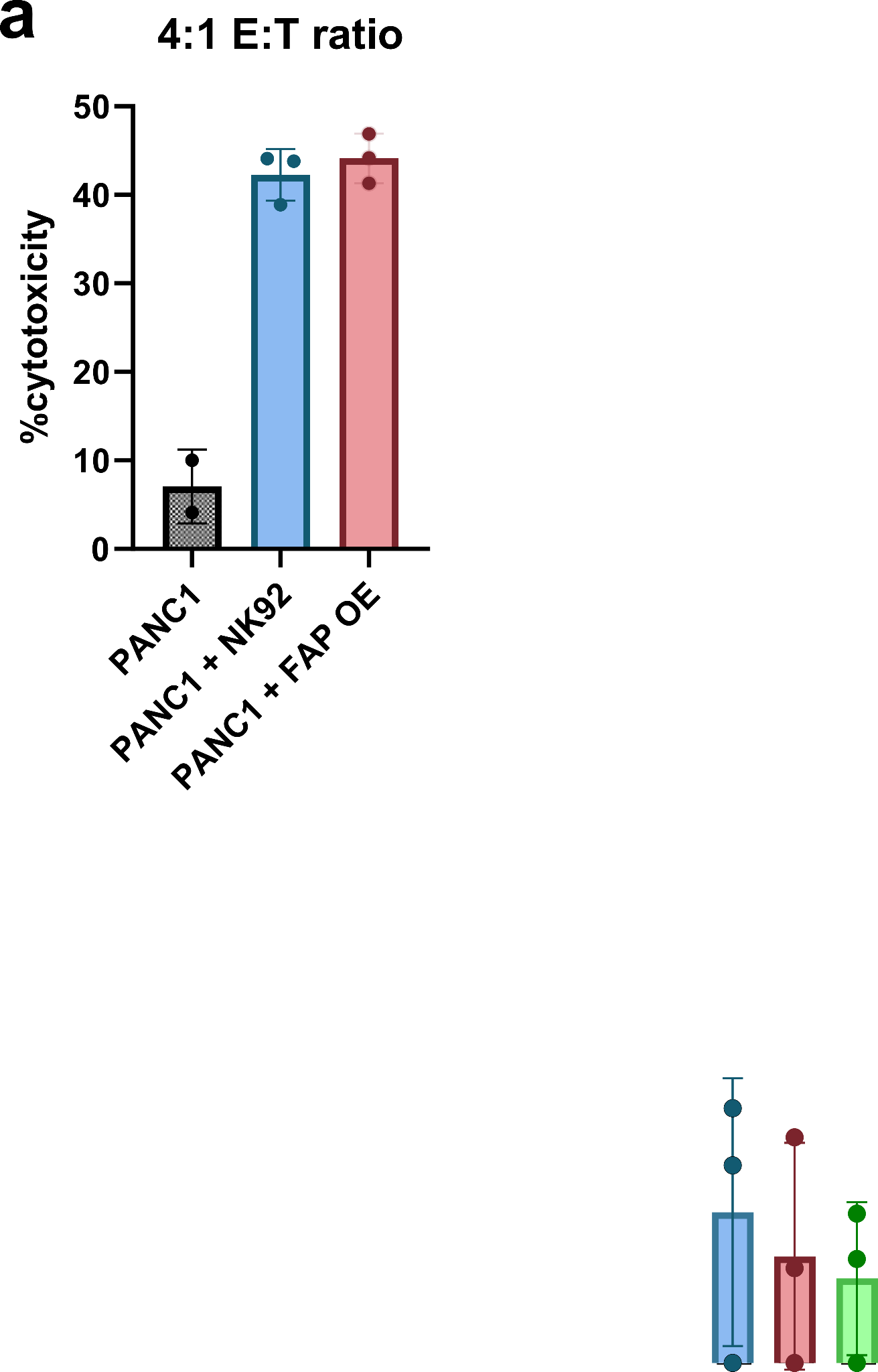

**Supplementary Fig. 9.** Effects of FAP manipulation on NK92 cytotoxicity. (**a**) Impact of FAP

OE on NK92 lysis of PANC-1 cells in 2D culture as determined by annexin flow cytometry assay. The effector to target ratio of NK to PANC-1 was 4 to 1. Cells were cocultured for 4 hours. Data is combined from three separate donors. All plots represent mean ± SD. two-tailed t-test. All bar plots represent mean ± SD.

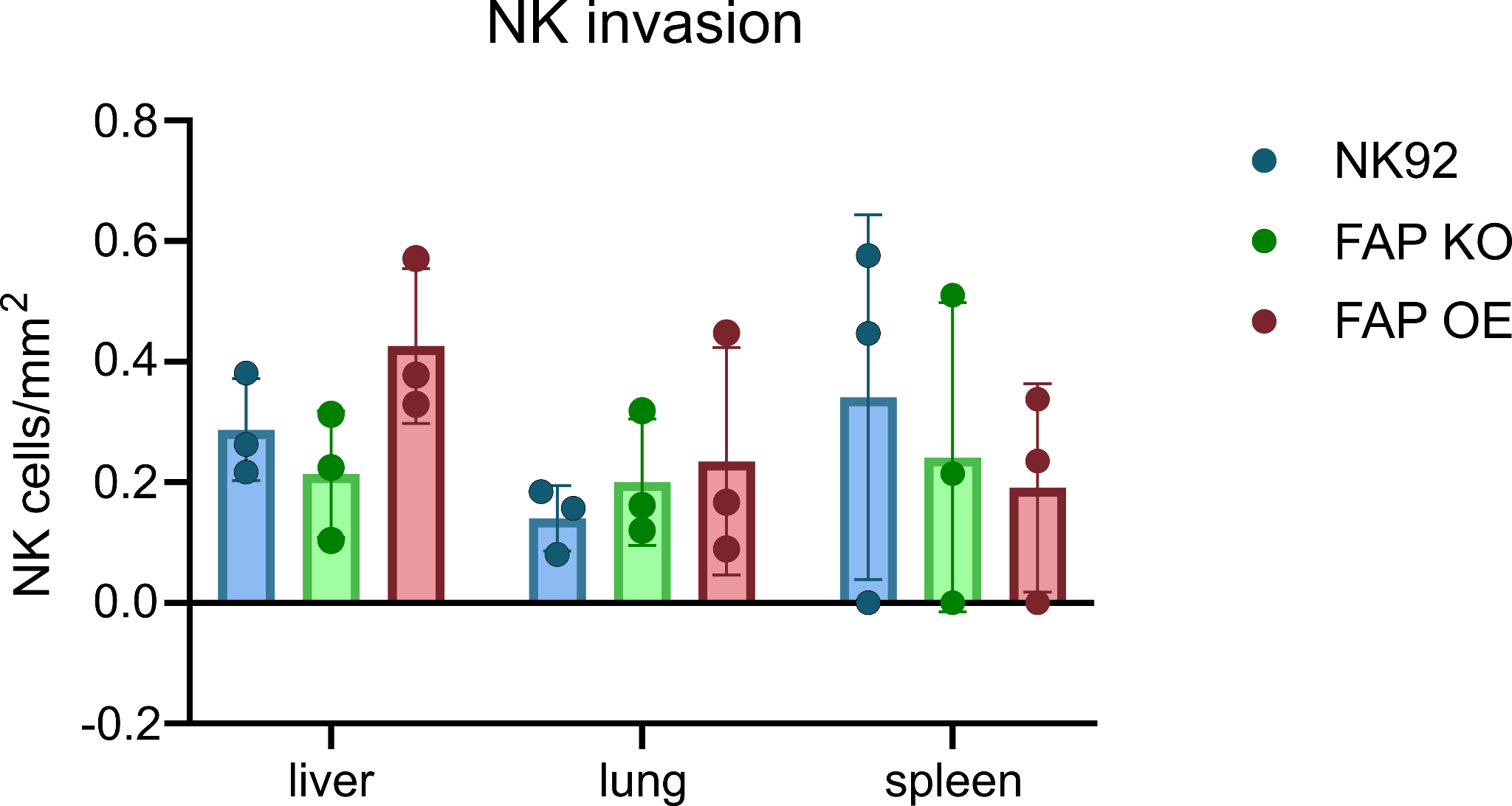

**Supplementary Fig. 10.** NK cell infiltration into organs. (**a**) Quantification of NK92, FAP OE NK92, and FAP KO NK92 cells per mm^2^ in organs. n=3 mice. All bar plots represent mean ± SD. No p-values were significant by unpaired two-tailed t-test.
